# The First High-Quality Reference Genome of Sika Deer Provides Insights for High-Tannin Adaptation

**DOI:** 10.1101/2021.05.13.443962

**Authors:** Xiumei Xing, Cheng Ai, Tianjiao Wang, Yang Li, Huitao Liu, Pengfei Hu, Guiwu Wang, Huamiao Liu, Hongliang Wang, Ranran Zhang, Junjun Zheng, Xiaobo Wang, Lei Wang, Yuxiao Chang, Qian Qian, Jinghua Yu, Lixin Tang, Shigang Wu, Xiujuan Shao, Alun Li, Peng Cui, Wei Zhan, Sheng Zhao, Zhichao Wu, Xiqun Shao, Yimeng Dong, Min Rong, Yihong Tan, Xuezhe Cui, Shuzhuo Chang, Xingchao Song, Tongao Yang, Limin Sun, Yan Ju, Pei Zhao, Huanhuan Fan, Ying Liu, Xinhui Wang, Wanyun Yang, Min Yang, Tao Wei, Shanshan Song, Jiaping Xu, Zhigang Yue, Qiqi Liang, Chunyi Li, Jue Ruan, Fuhe Yang

## Abstract

Sika deer are known to prefer oak leaves, which are rich in tannins and toxic to most mammals; however, the genetic mechanisms underlying their unique ability to adapt to living in the jungle are still unclear. In identifying the mechanism responsible for the tolerance of a highly toxic diet, we have made a major advancement in the elucidation of the genomics of sika deer. We generated the first high-quality, chromosome-level genome assembly of sika deer and measured the correlation between tannin intake and RNA expression in 15 tissues through 180 experiments. Comparative genome analyses showed that the *UGT* and *CYP* gene families are functionally involved in the adaptation of sika deer to high-tannin food, especially the expansion of *UGT* genes in a subfamily. The first chromosome-level assembly and genetic characterization of the tolerance toa highly toxic diet suggest that the sika deer genome will serve as an essential resource for understanding evolutionary events and tannin adaptation. Our study provides a paradigm of comparative expressive genomics that can be applied to the study of unique biological features in non-model animals.

## Introduction

Cervidae consists of 55 extant deer species and constitutes the second largest family in terrestrial artiodactyls. Sika deer (*Cervus nippon*) is naturally distributed throughout East Asia and is one of the best-known deer species producing velvet antlers [1,2], a valuable ingredient in traditional Chinese medicine [3]. Among other deer species [4–6], sika deer has unique characteristics, such as a geographic distribution that is significantly more coincident with oak trees (Figure 1A) and an ability to tolerate a high-tannin diet, mainly consisting of oak leaves. Notably, oak leaves, which are rich in tannins and toxic to most mammals, such as cattle, which are related to sika deer [7], are conversely found to increase the reproductive rate and fawn survival rate of sika deer. Thus, oak leaves are essential for maintaining healthy sika deer in wild and farmed populations. Some studies have concluded that tannins are not toxic to sika deer because of the rumen microbes and fermentation patterns of these deer [8]. However, knowledge is scarce regarding the genetics and mechanism underlying the ability to detoxify a high-tannin diet.

**Figure 1.**
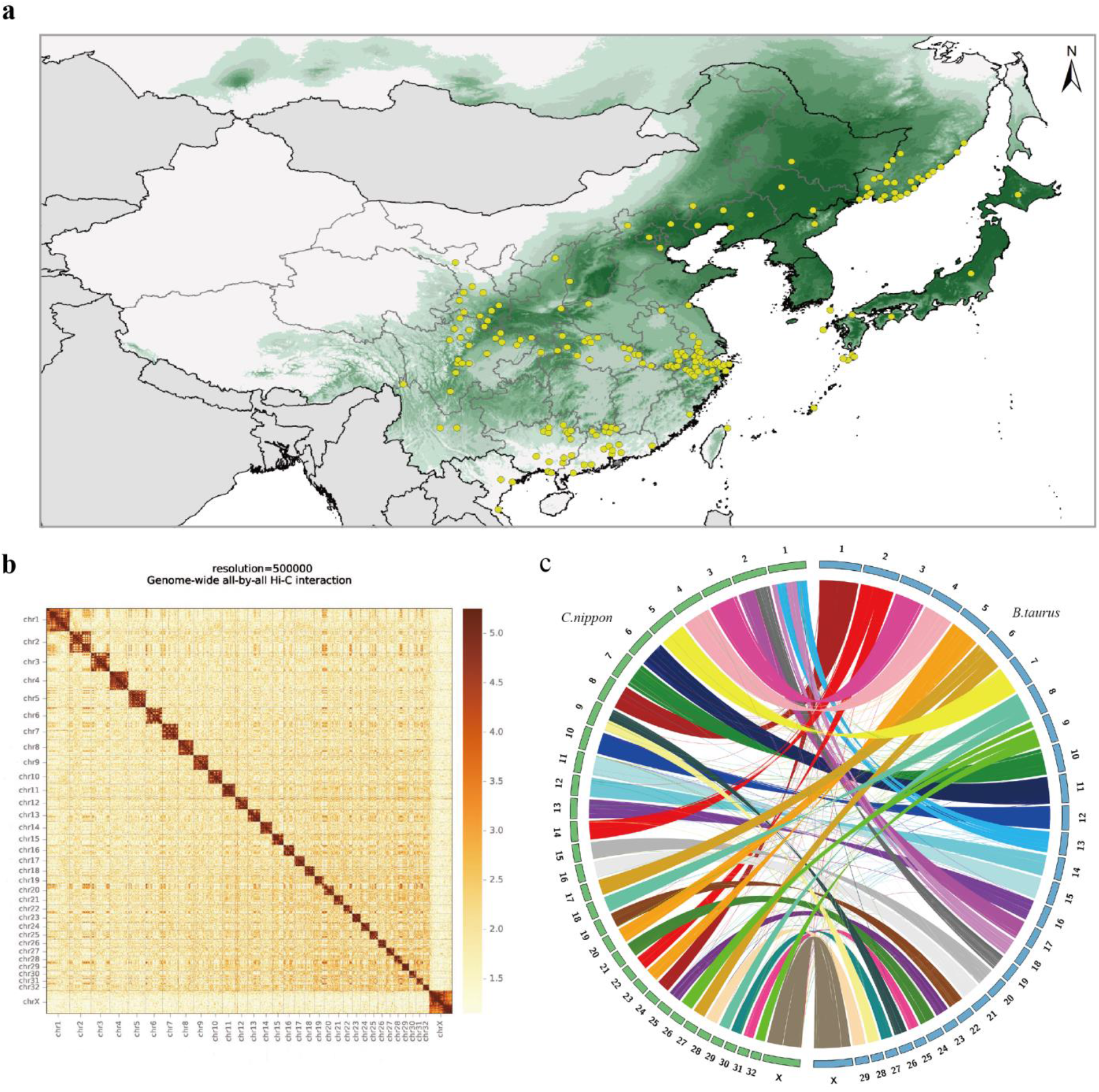
Distribution and genome assembly of sika deer. **A**, Mongolian oak and sika deer distribution. The green shadow represents the distribution range of Mongolian oak. The yellow dots represent the historical distribution of sika deer in 5 countries (China, Russia, Japan, North Korea and Vietnam). **B**, A contact map at a 500-kb resolution of chromosome-level assembly in sika deer is shown. The color bar illuminates the logarithm of the contact density from red (high) to white (low) in the plot. Note that only sequences anchored on chromosomes are shown in the plot. **C**, Synteny analysis of cattle and sika deer. Circular graphs displaying the results from the synteny analysis. Same-color ribbons connect syntenic genomic segments.

Whole-genome sequencing has become a more popular technology with which to explore the taxonomy, evolution and biological phenomena of organisms at the molecular level [9], compared with morphological, histological and other analyses [10–12]. For example, a series of studies investigated the genomes of 11 deer and 33 other ruminant species and identified some genes that are involved in ruminant headgear formation, rapid antler regeneration, and reindeer adaptation to the long days and nights in the Arctic region [6,13,14]. The chromosome-level reference genome for sika deer is in high demand compared with that for other ruminants such as bovines [15,16], and it will provide novel genomic and molecular evolutionary information on the exceptional characteristics of the sika deer.

Here, we report the chromosome-level genome assembly of a female sika deer, as well as the RNA sequencing of 15 tissue types in sika deer treated with 3 levels of a high-tannin diet. The findings provide important resources to help elucidate the genetic mechanisms underlying the high-tannin food tolerance of sika deer. Our high-quality sika deer genome will be of great importance to researchers who study the common characteristics of deer and other ruminants and could even serve as a reference deer genome. The well-designed RNA expression experiments used in this study also provide a paradigm for studying novel features in nonmodel animals.

## Results

### De novo assembly of a Cervus nippon reference genome

We collected DNA from a female sika deer (*Cervus nippon*) and identified a total of 66 chromosomes, including 64 autosomes and one pair of sex chromosomes (XX) (Additional file 1: Figure S1). A large set of data was acquired for assembly using a combination of four technologies. 1) A total of 242.9 Gb of clean data (~93.4×) were obtained from paired-end sequencing (Illumina HiSeq), with the genome size (2.6 Gb) estimated by the 25 K-mer distribution (Additional file 2: Table S1 and Additional file 1: Figure S2). 2) A total of 150.4 Gb (~57.7×) of PacBio RSII long reads (single-molecule real-time sequencing) were also acquired (Additional file 2: Table S2). The wtdbg2 [17] assembler yielded 2,040 primary contigs using PacBio reads with a contig N50 size of 23.6 Mb and the longest at 93.6 Mb (Additional file 2: Table S3). These contigs were then polished using the Quiver algorithm [18] with default parameters. Genome-wide base-level correction was performed using Illumina short reads aligned to the published genome with BWA (v0.7.10-r943-dirty), and inconsistencies between the genome and the reads were identified with SAMTools/VCFtools (v1.3.1). These inconsistencies were corrected by our in-house script to produce a highly accurate assembly. 3) The previous contigs were clustered into chromosome-scale scaffolds using high-throughput chromosome conformation capture (Hi-C) proximity-guided assembly (Figure 1B) to produce the final reference assembly, named MHL_v1.0, totaling 2.5 Gb of sequence with a contig N50 of 23.6 Mb and a scaffold N50 of 78.8 Mb (**Table 1**). The resulting assembly contained 2,481,763,803 bp reliably anchored on chromosomes, accounting for 99.24% of the whole genome (Additional file 2: Table S4). 4) A total of 264 Gb of optical mapping (using BioNano Genomics Irys) data were also used to generate *de novo*-assembled optical maps with a scaffold N50 of 1.974 Mb, which was sequentially compared with MHL_v1.0 to identify the misoriented contigs and improve the final validated reference assembly (Additional file 1: Figure S3).

**Table 1.**
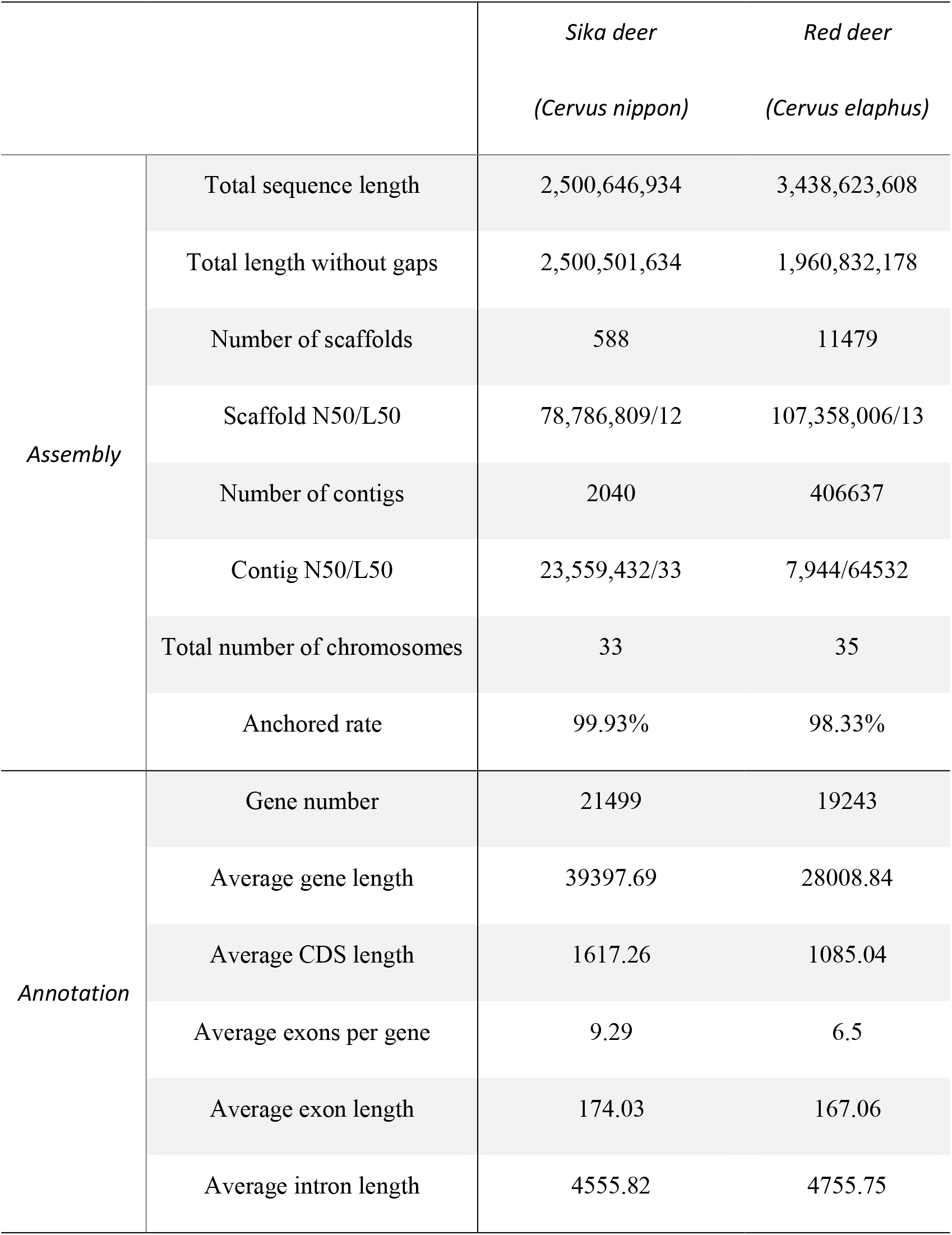
Comparison of genome quality and annotation between the genome of sika deer and the best published genome of red deer

|  |  | *Sika deer*<br>*(Cervus nippon)* | *Red deer*<br>*(Cervus elaphus)* |
| --- | --- | --- | --- |
| *Assembly* | Total sequence length | 2,500,646,934 | 3,438,623,608 |
|  | Total length without gaps | 2,500,501,634 | 1,960,832,178 |
|  | Number of scaffolds | 588 | 11479 |
|  | Scaffold N50/L50 | 78,786,809/12 | 107,358,006/13 |
|  | Number of contigs | 2040 | 406637 |
|  | Contig N50/L50 | 23,559,432/33 | 7,944/64532 |
|  | Total number of chromosomes | 33 | 35 |
|  | Anchored rate | 99.93% | 98.33% |
| *Annotation* | Gene number | 21499 | 19243 |
|  | Average gene length | 39397.69 | 28008.84 |
|  | Average CDS length | 1617.26 | 1085.04 |
|  | Average exons per gene | 9.29 | 6.5 |
|  | Average exon length | 174.03 | 167.06 |
|  | Average intron length | 4555.82 | 4755.75 |

To validate our assembly, MHL_v1.0 was compared with the previously published red deer [19] genome (Additional file 1: Figure S4). Both the inconsistency of the synteny analysis and the improper density of Hi-C proximity maps identified 34 inaccurate junctions, which were considered potential inversions and misassemblies (Additional file 1: Figures S4 and S5). The aforementioned optical maps were used to determine whether the 34 inaccurate junctions were breakpoints or new joint regions after the replacement. We found that 10 inaccurate junctions were supported by the optical maps, and those junctions were then manually inspected and correlated. Additionally, another 142 potential misjoined contigs were found by comparing our MHL_v1.0 assembly with the optical maps. The paired-end Illumina short reads were then mapped to the final assembly, and all 142 disagreements were checked manually and found to be sequential in the comparison results. We further compared MHL_v1.0 with the twenty published genomes of Cervidae, including red deer (*Cervus elaphus*) and reindeer (*Rangifer tarandus*). The results showed that the scaffold N50 length and ungapped sequence length of the MHL_v1.0 assembly were greater than those previously published (Additional file 2: Table S5). We compared three other chromosome-level ruminant genomes (cattle, goat, and red deer) with MHL_v1.0. Multiple chromosome fission/separation events were detected among these four genomes, and we found that the sika deer genome had the highest chromosome collinearity with red deer (Figure 1C and Additional file 1: Figure S6).

Finally, we downloaded a total of 2,715 EST sequences belonging to sika deer from the NCBI dbEST database and aligned them against MHL_v1.0. We found that 95.95% of the EST sequences (coverage rate > 90%) matched our sika deer genome MHL_v1.0. Evaluation of our MHL_v1.0 using CEGMA software showed that 97.18% of the full length of 248 genes in the core gene set was predicted. Benchmarking Universal Single-Copy Orthologs (BUSCO) analysis of the gene set showed that complete BUSCO accounted for 3,880 (of 4,104; 94.60%) genes, which is better than the results obtained for the water buffalo (*Bubalus bubalis*, 93.6%) [12] and domestic goat (*Capra hircus, 82%*) [20]. After aligning Illumina short reads (93.4×) against MHL_v1.0, the base level error rate was estimated to be 1.1e-5 (Additional file 2: Table S6).

### Genome annotation

Homology and *de novo* repetitive sequence annotation results showed that repetitive sequences accounted for approximately 45.38% of MHL_v1.0, which is consistent with the percentages published for other mammals (Additional file 2: Tables S7 and S8), including humans (44.8%) [21], water buffalo (45.33%) [12] and sheep (42.67%) [22]. As in other published mammalian genomes, long interspersed nuclear elements (LINEs), short interspersed nuclear elements (SINEs) and long terminal repeats (LTRs) were also the most abundant elements in MHL_v1.0 (29.56%, 7.63% and 5.38% of the total number of elements, respectively) (Additional file 1: Figure S7). The main features of MHL_v1.0 are summarized and shown in Additional file 1: Figure S8.

A total of 21,449 protein-coding genes were predicted using the combined methods of homology and *de novo* annotations with transcriptome data (mapping rate of 93.43% for 1.2 billion RNA-Seq reads), and 90.1% of the protein-coding genes were functionally annotated (Additional file 2: Table S9). The average coding sequence (CDS) length per gene was 1,617 bp, the exon number per gene was 9.29, and the average length per exon was 174 bp; these values are similar to those in other mammals (Additional file 2: Table S10). To verify the accuracy of our gene predictions and to assess the annotation completeness of MHL_v1.0, we checked core gene statistics using the BUSCO software. A total of 3,907 (of 4,104; 95.20%) (Additional file 2: Table S11) highly conserved core proteins in mammals were recovered from our predictions.

### Analyses of phylogeny and demographics

A phylogenetic tree (**Figure 2A**) based on 19 mammals spanning the orders Primates, Rodentia, Artiodactyla and Cetacea was constructed with the maximum-likelihood method using 748 identified single-copy orthologous genes. The results showed that sika deer was in the same clade as red deer (Figure 2A), which is consistent with the cladistic data [23]. The divergence time between sika deer and red deer was estimated to be approximately 2.5 million years ago (MYA) (Figure 2A and Additional file 1: Figure S9).

**Figure 2.**
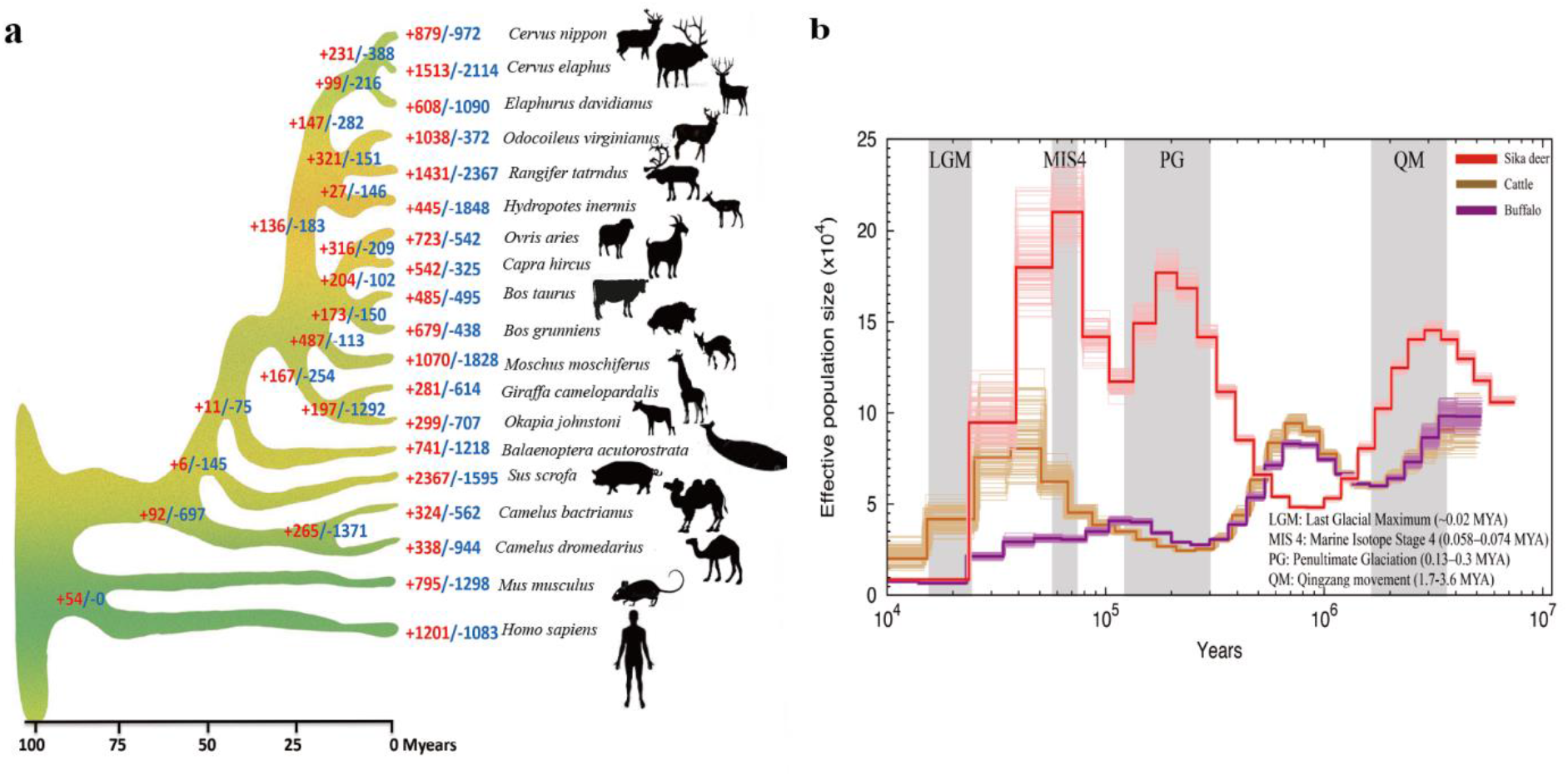
Evolutionary analysis of sika deer. **A**, Phylogenetic tree inferred from 19 species. The x-axis is the inferred divergence time (M years) based on the phylogenetic tree and fossils. The number of expanded gene families is red, and the number of contracted gene families is blue. **B**, PSMC analysis of effective population sizes in sika deer, cattle and buffalo.

To examine the changes in effective population size (Ne) of the ancestral populations, a Pairwise Sequential Markovian Coalescent (PSMC) analysis was applied to sika deer, cattle [16] and buffalo [12] (Figure 2B). Demographic analysis showed that the Ne of the sika deer sharply declined during the two large glaciations: the Qingzang movement (1.7-3.6 MYA) and Penultimate Glaciation (0.13-0.3 MYA), and the sika deer underwent a long period of population bottlenecks. Subsequently, the Ne increased greatly after that period, suggesting that these deer had adapted to the specific habitat, probably due to the monsoon climate in East Asia. During the same period, the populations of cattle and buffalo recovered soon after a decline and shrank again. During Marine Isotope Stage 4 (0.058-0.074 MYA) and the last glacial maximum (LGM, ~0.02 MYA), sika deer suffered population bottlenecks again (Figure 2B), which may also be the reason modern sika deer populations have very low genetic diversity [23].

### Gene family evolution

We identified a total of 9,830 homologous gene families in MHL_v1.0 by comparing the predicted protein sequences of sika deer with those of 19 mammals spanning the orders Primates, Rodentia, Artiodactyla and Cetacea (Additional file 2: Table S12 and Additional file 1: Figure S10).

Based on the hypothesis that potential genomic adaptations are related to genes that are under positive selection in the sika deer lineages [24], we identified 55 positively selected genes (PSGs), which were calculated using the branch-site models and validated using likelihood ratio tests (Additional file 2: Table S13). The PSGs were found to be involved in the PI3K-Akt signaling pathway (ko04151), VEGF signaling pathway (ko04370) and pathways in cancer (ko05200), among others. These pathways were reportedly related to antler growth [25,26].

The number of genes in a gene family has been proposed as a major factor underlying the adaptive divergence of closely related species. To depict the gene family evolution, we identified 972 significantly contracted and 879 significantly expanded gene families in sika deer compared with other species (Figure 2A). The expanded gene families were mainly enriched in the signal transduction pathways of environmental perception (olfactory transduction, G protein-coupled receptors, neuroactive ligand-receptor interaction, corrected *P*-value < 0.05), enzymatic activity (transferase activity, transferring hexosyl groups, carboxypeptidase activity and L-lactate dehydrogenase activity, corrected *P*-value < 0.05), feeding behavior (salivary secretion, neurotransmitter secretion, corrected *P*-value < 0.05) and drug metabolism (drug metabolism - other enzymes, drug metabolism - cytochrome P450, metabolism of xenobiotics by cytochrome P450, corrected *P*-value < 0.05) (Additional file 2: Tables S14 and S15). The contracted gene families were mainly related to lipid metabolism pathways (linoleic acid metabolism and ether lipid metabolism, corrected *P*-value < 0.05), ion transportation (calcium ion binding, anion transport, and iron ion binding, corrected *P*-value < 0.05) and regulation of basic biological processes (regulation of developmental and apoptotic processes, corrected *P*-value < 0.05) (Additional file 2: Tables S16 and S17).

### Exceptional expansion of the UGT gene family in the sika deer genome

Gene gains and losses are one of the primary contributors to functional changes. To better understand the evolutionary dynamics of genes, we assessed the expansion and contraction of the gene ortholog clusters among 19 species. The uridine 5’-diphospho-glucuronosyltransferase (UDP-glucuronosyltransferase, *UGT*) gene families were at the top 27 of 879 significantly expanded gene families, which have been reported to play a role in the catabolism of exogenous compounds [27–29]. Phylogenetic analysis revealed that the 257 *UGT* genes could be classified into 7 lineages (Figure 3A and Additional file 1: Figure S11), while in the sika deer genome, we found two lineage-specific monophyletic expansions of the *UGT2B* and *UGT2C* subfamilies (Figure 3B). In the *UGT2B* subfamily, 15 copies were found in the sika deer genome, which was more than that in any other species assessed in this study (Additional file 2: Table S18). Sika deer had relatively lower expanded gene numbers in the *UGT2C* subfamily than in the *UGT2B* subfamily (Additional file 2: Table S18). Taken together, these results prompt us to propose that the exceptional expansion of the *UGT* gene family may be the key genetic basis for the tolerance of high-tannin food, namely, oak leaves, by the sika deer.

**Figure 3.**
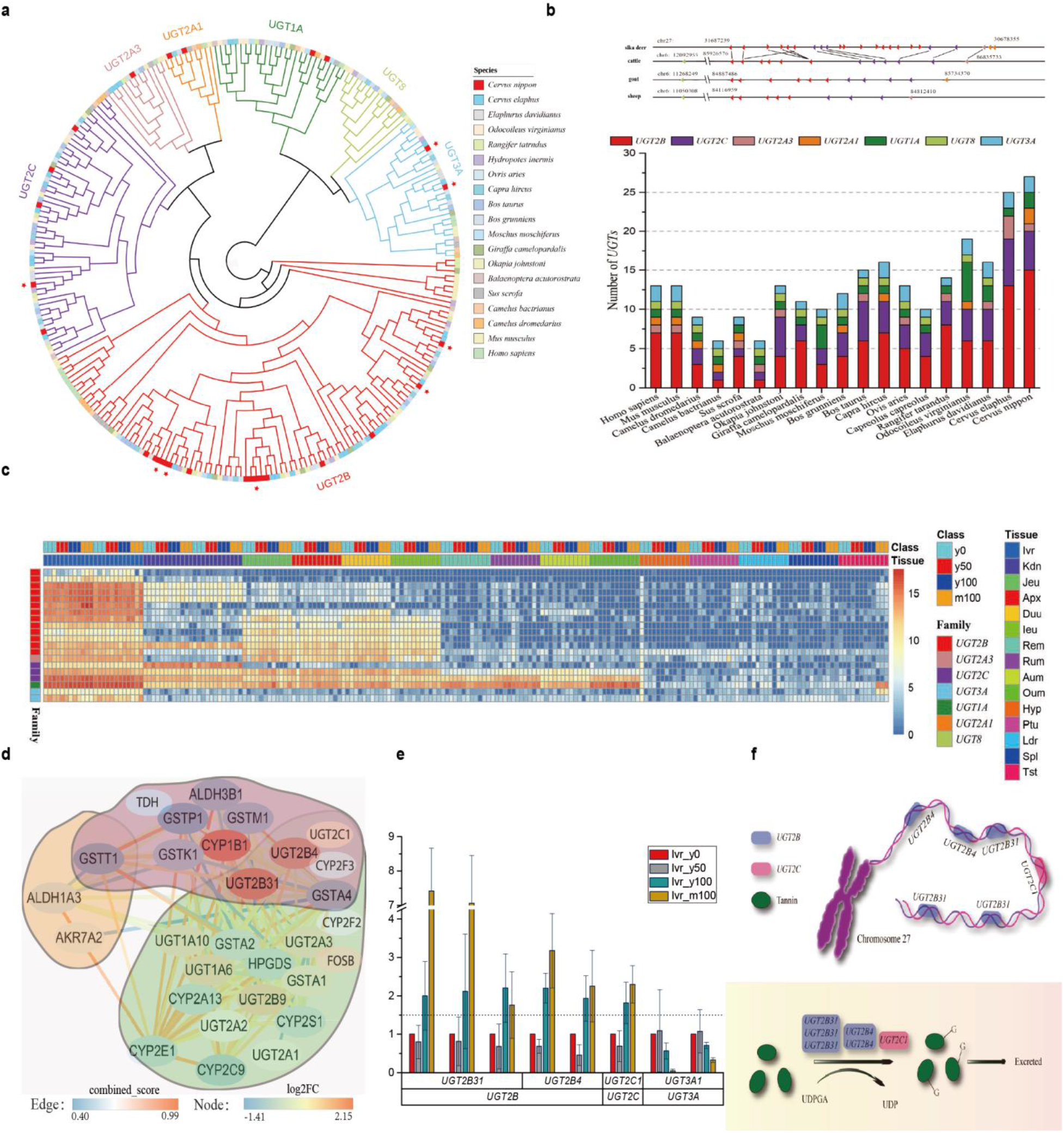
*UGT* expansion and high-tannin adaptation in sika deer. Transcriptome analysis revealed that the *UGT* gene family was the key factor for sika deer adaptation to a high-tannin diet. **A**, Gene tree of *UGTs* in 19 species. The red stars are significantly differentially expressed genes in the sika deer transcriptome. **B**, Number of *UGT* genes in 19 species. **C**, Expression heatmap of *UGTs* of sika deer in different tissues and treatments. **D**, The overlap between 3 contracted genes (yellow background), 20 expanded genes (green background) and 12 DEGs (pink background), which all play a role in the cytochrome P450 pathway. **E**, Expression change of 8 significant differentially expressed genes in sika deer liver resulting from different treatments. **F**, Six upregulated *UGT* genes in the *UGT2B* and *UGT2C* subfamilies were located on sika deer chromosome 27; schematic of the glucuronidation reaction. UDPGA, uridine 5’-diphospho-glucuronic acid.

### Transcriptomic analysis of 15 tissues of sika deer treated with a high-tannin diet

Sika deer adapted well to living in the forest and have consumed a high-tannin diet of Mongolian oak (*Quercus mongolica*) leaves (MOL) for a long time; whether the underlying genetic adaptation and molecular mechanism are associated with the special expansion of *UGT* gene families is an interesting question. We used 9 deer fawns to conduct a feeding trial with different tannin-containing (0%, 50%, 100%) diets, and 3 mature deer (100%) were used as a comparison group. Transcriptome sequencing was performed on 15 tissues of all experimental individuals (Additional file 2: Table S19). A total of 1.44 Tb of transcriptional data from 180 samples were obtained using the Illumina platform, and the 17,233 differentially expressed genes (DEGs) were analyzed by pairwise comparison of each group (Additional file 1: Figure S12). The liver is the major organ associated with *UGT* activity, and *UGT* expression was highest in the liver among the fifteen tissues examined (Figure 3C). Although *UGT* genes were also highly expressed in the liver tissue of cattle, they did not respond to high MOL levels (Additional file 1: Figure S13). We compared different MOL levels in sika deer and identified 3,222 and 15 DEGs in liver and kidney tissue, respectively.

After inspecting all the expanded/contracted gene families and DEGs in liver tissue, 29 genes were found to play roles in the P450 pathway. Of these, 20 were expanded genes, 12 were DEGs, and 3 were contracted genes. The interaction network of these genes is shown in Figure 3D. Among these key genes, *UGT2B4* and *UGT2B31* were both significantly upregulated in high-tannin liver tissue and expanded in the sika deer genome. Therefore, we hypothesized that *UGT2B4* and *UGT2B31* are major genes in sika deer with high-tannin adaptation.

Interestingly, in liver tissue, tannins can drive the expression of many *UGT* genes in a dose-dependent manner. Overall, when compared among different MOL levels and ages (y0, y50, y100 and m100), eight differentially expressed *UGT* genes were discovered, among which two were downregulated genes of the *UGT3A* subfamily and six were upregulated genes in the *UGT2B* and *UGT2C* subfamilies (Figure 3E). Furthermore, we found that all of these upregulated *UGT* genes in the liver were located on sika deer chromosome 27 (Figure 3F). With the increase in tannin content intake, the *UGT3A* subfamily genes in the liver were inhibited; nevertheless, *UGT* gene copies in the *UGT2B* and *UGT2C* families were increased, suggesting that the response of *UGT* gene expression to tannin was mainly upregulated. Moreover, in the kidney tissue, two DEGs belonged to the *UGT2C* family. Five differentially expressed *CYP* genes were upregulated, whereas gene families encoding *GST* and *SULT* were all downregulated after the deer were fed a high-tannin diet. According to previous studies, sika deer share common pathways with koala, including the drug metabolism-cytochrome P450 signal pathway [11]. The detoxification genes in sika deer showed opposite expression patterns compared with the genes in koala [11] (Additional file 1: Figure S14). These results indicate that sika deer may utilize a different adaptive strategy from that of koala to survive on a diet of highly toxic food.

### Ability to tolerate a high-tannin diet

The sika deer diet of MOL contains high levels of tannins that would be lethal to most other mammals. The main detoxification reactions are traditionally categorized into phase I and phase II reactions. Currently available evidence indicates that among these, the *CYP*, *UGT*, *GST*, and *SULT* gene families have the greatest importance in xenobiotic metabolism. Based on the aforementioned mechanism, genes involved in those pathways were examined using gene family and transcriptome analyses.

A total of 13 DEGs were detected from the *CYP2* family in sika deer liver, but only 5 were differentially expressed with increasing tannin contents in the diet. Five *GST* genes and 3 *SULT* genes were found to be differentially expressed in the liver, but all were downregulated with increasing tannin contents in the diet.

The functional importance of these *UGT* genes was further investigated through analysis of their expression levels in sika deer, showing that they had particularly high expression in the liver tissue, which is consistent with their role in detoxification. The mechanism of the glucuronidation reaction is that *UGT* enzymes catalyze the transfer of the glucuronosyl group from uridine 5’-diphospho-glucuronic acid (UDPGA) to the tannin molecules, generating the glucuronidated metabolite, which is more polar and more easily excreted than the tannin molecule (Figure 3F). Most of these expressed *UGT* genes belonged to *UGT2B*. These phenotypes suggest that *UGT* genes in *UGT2B* have an important role in detoxification; the upregulated expansion of *UGT* genes would result in higher enzyme levels, which would enhance the ability of sika deer to detoxify the high-tannin diet.

Among the genes related to the metabolism of drugs and exogenous substances, *UGT* and *CYP* genes were found to be functionally involved in detoxification, especially *UGT* genes in the *UGT2B* family. In short, these findings imply that the unique expansion of the *UGT* gene family is mainly responsible for the toleration of high-tannin food, namely, oak leaves, by sika deer (Additional file 1: Figure S15).

## Discussion

Cervidae is the second largest family in Artiodactyla [30] and has significant scientific [1] and economic [3] value. Although several other deer genomes have recently been reported, the lack of high-quality genome sequences of sika deer, one of the novel species in the family, has hindered the elucidation of the molecular mechanisms underlying important distinct biological characteristics of sika deer, such as the full regeneration of the antlers. Here, we sequenced the genome of sika deer and assembled it at the chromosome level using combined technologies of SMRT, Illumina sequencing and Hi-C. The high percentage and accuracy rate of the genome structure, base calling, gene set validation and quality of gene annotation demonstrated that our assembled sika deer genome was of high quality and could be effectively used as a reference genome for deer species.

The geographic distribution of sika deer is highly coincident with that of oak, and sika deer have a preference for grazing on high-tannin oak leaves [31], suggesting that this adaptation may be a positive selection during evolution. In terms of food adaption, sika deer are not unique. For example, pandas, dogs and koalas have also undergone adaptive food evolution; pandas can eat bamboo despite being carnivorous [32], dogs can adapt to a diet of starchy foods [33], and koalas can eat toxic eucalyptus leaves [11]. Divergent adaptive pathways and related genes are known to be involved in this adaptation. In this study, we found that among the genes related to toxin degradation, only those from the *UGT* gene family [34], especially the *UGT2B* family, were significantly expanded. Furthermore, transcriptomic studies showed that *UGT* gene expression was strongly correlated with the quantity of tannin intake, i.e., it was dose dependent. The expression of specific extended gene copies in the *UGT2B* family was prominently increased after the tannin feeding treatment. These results suggest that genes in the *UGT* family, especially in the *UGT2B* subfamily, are associated with the adaptation of sika deer to a high-tannin diet.

It is generally believed that rumen microorganisms play a role in the digestion of tannins [35,36]. However, as other ruminants, such as cattle and sheep, are not well adapted to high-tannin diets (Additional file 1: Figure S16), we speculate that during a long period of coexistence with oak trees during evolution, sika deer may have developed genetic adaptive mechanisms. As expected, we found evidence for this phenomenon at the genome level through high-quality sequencing. Transcriptomic results also revealed that changes in gene expression were involved in Na and K ion channels. The Na and K balance (water and salt metabolism) is essential for the basic metabolism of organisms. These genetic responses have enabled sika deer to adapt to oak leaves as an advantageous rather than a hazardous material for consumption.

## Conclusion

The sika deer genome assembled in this study provides, to our knowledge, the highest quality deer genome to date. The comprehensive characterization of the sika deer genome along with the transcriptomic data presented herein provides a framework used to elucidate its evolutionary events, revealing the mechanism of the unique attributes and tannin adaptation. Through detailed genomics and transcriptomics analyses, we identified the most likely mechanism of tannin degradation in sika deer. We also depicted possible molecular mechanisms for the jungle adaptability of deer, and the methodologies we used in this study will also provide a reference for the study of the adaptation mechanism of animals to “toxic” foods. Chromosome-scale assembly of sika deer genomes could be used for many applications, including the study of structural variations in large genomic regions, expected recombination frequencies in specific genomic regions, target sequence characterization and modification for gene editing. Moreover, this study provides a valuable genomic resource for research on the genetic basis of sika deer’s distinctive physiological features, such as the full regeneration of deer antlers, and on Cervidae genome evolution. Our study also contributes to conservation and utilization efforts for this antler-growing species.

## Materials and methods

### Method details

#### Sampling preparation

A female sika deer (*Cervus nippon*) from Jilin Province was used for *de novo* genome sequencing. DNA was extracted from whole blood with a BioTeke DP1102 kit (solution) according to the manufacturer’s instructions. After slaughtering the experimental animals, tissue sampling was carried out immediately. Tissues, such as those from the hypothalamus, pituitary, gonad, liver, kidney, spleen, rumen, reticulum, and small intestine, were collected. RNA was extracted from the 15 tissue samples obtained from the animals. After library construction and size selection, 150.4 Gb (57.7×) of long reads with a mean length of 9,205 bp were generated by the PacBio RSII platform (Single Molecule Real-Time, SMRT). In addition, 261.5 Gb (100.6×) of paired-end data with varying insert sizes (200, 300, 400, and 600 bp) were generated by the Illumina HiSeq 2000 platform (Additional file 1: Figure S17).

#### De novo *genome sequencing and Hi-C-based assembly*

The PacBio subreads were used to perform *de novo* genome assembly via wtdbg2 [17] with the key parameter “-H –k 19”. Then, primary assemblies were polished using the Quiver [18] algorithm with the default parameters. A total of 93.4× clean paired-end reads from the Illumina platform were aligned to the Quiver-polished assemblies using BWA (v0.7.10-r943-dirty) to reduce the remaining InDel and base substitution errors in the draft assembly. Inconsistent sequences between the polished genome and Illumina reads were identified with SAMTools/VCFtools (v1.3.1). The credible homozygous variations with differences in quality exceeding 20, a mapping quality greater than 40 and a sum of high-quality alt-forward and alt-reverse bases more than 2 in the Quiver-polished assemblies were replaced by the called bases using in-house scripts. Finally, highly accurate contigs were generated.

Four billion PE150 reads were produced from three Hi-C libraries by the Illumina HiSeq platform. Hi-C-based proximity-guided scaffolding was used to connect primary contigs. Clean reads were first aligned against the reference genome with the Bowtie2 end-to-end algorithm. HiC-Pro (v2.7.8) was then able to detect the ligation sites and align them back to the genome with the 5’ fraction of the reads. The assembly tool LACHESIS was applied for clustering, ordering and orienting. Based on the agglomerative hierarchical clustering algorithm, we clustered the contigs into 33 groups. For each chromosome cluster, we obtained an exact scaffold order of the internal groups and traversed all the directions of the scaffolds through a weighted directed acyclic graph (WDAG) to predict the orientation for each scaffold. A chromosome-scale assembly with 33 clusters was obtained that anchored 99.24% of the contigs for sika deer.

#### Genome accuracy assessment

To determine the completeness and accuracy of the MHL_v1.0 assembly, we carried out the following validation. First, the MHL_v1.0 assembly was aligned to the red deer genome (CerEla1.0) and BioNano optical maps. The conflicting regions that appeared in both alignments were potential misassemblies and were manually inspected andcorrected.

A total of 2,715 EST sequences of sika deer were downloaded from the NCBI dbEST database and aligned with MHL_v1.0 using BLAST (v35). The BUSCO [37] software package was used to assess the quality of the generated genome using the genome model M genome”. The CEGMA pipeline software, which was also run against the MHL_v1.0. Illumina short reads (93.4×), was aligned to MHL_v1.0 with BWA to estimate the accuracy of a single base of the assembly, which was based on the count of homozygous SNPs.

#### Repeat sequence annotation

To annotate the sika deer genome, RepeatModeler (v1.0.8) was initially used to obtain a *de novo* repeat library. Next, RepeatMasker (v4.0.5) was used to search for known and novel transposable elements (TEs) by mapping sequences against the Repbase TE library (20150807) [38].

#### Gene annotation

For *de novo* gene prediction, we utilized AUGUSTUS (v3.0.3), SNAP (v2006-07-28), GlimmerHMM (v3.0.4) and GENSCAN to analyze the repeat-masked genome. For homology-based gene predictions, the protein sequences of human, mouse, cattle, sheep, and horse were mapped to the sika deer genome with GenBlastA [39]. Then, prediction was performed with GeneWise (v2.2.3) [40] in aligned regions. RNA-seq reads were aligned to the genome using TopHat (v2.0.12) and assembled by Cufflinks (v 2.2.1) with the default parameters. EVidenceModeler software (EVM, v1.1.1) was used to integrate the genes predicted by homology, *de novo* and transcriptome approaches and generate a consensus gene set. Short-length (< 50 aa) and transcriptome data for nonsupport genes were removed from the consensus gene set, and the final gene set was produced.

We translated the final predicted coding regions into protein sequences and mapped all the predicted proteins to the Swiss-Prot, TrEMBL, and KEGG databases using BLASTP (v2.2.27+) for gene functional annotation. We used the InterProScan database to annotate the motifs, domains and Gene Ontology (GO) terms of proteins with retrieval from the Pfam, PRINTS, PROSITE, ProDom, and SMART databases.

#### Gene family construction

Annotations of human, mouse, pig, sheep and cattle genomes were downloaded from Ensembl (release-87), while those of minke whale, dromedary, Bactrian camel, yak, goat, white-tailed deer, red deer, and reindeer were downloaded from NCBI. To annotate the structures and functions of putative genes in the giraffe, okapi, milu, musk deer, and roe deer assemblies, we used homology-based predictions. Cattle proteins (Ensemble release-87) were aligned to the 5 genomes using GenBlastA (v1.0.1) [39] and predicted by GeneWise (v2.2.3) [40]. The genes of the above 18 species and sika deer were used to construct gene families using TreeFam [17]. All the protein sequences were searched in the TreeFam (version 9) HMM file and classified among different TreeFamilies.

#### Phylogeny and divergence time estimation

We constructed a phylogenetic tree based on a concatenated sequence alignment of 748 single-copy gene families from sika deer and 18 other mammalian taxa (human, mouse, pig, sheep, cattle, minke whale, dromedary, Bactrian camel, yak, goat, white-tailed deer, red deer, reindeer, giraffe, okapi, milu, musk deer, and roe deer) using the RAxML [41] software with the GTRGAMMA model. Divergence times were estimated by PAML [42] MCMCTREE. The Markov chain Monte Carlo (MCMC) process was run for 20,000 iterations with a sample frequency of 2 after a burn-in of 1,000 iterations. Other parameters used the default settings of MCMCTREE. Two independent runs were performed to check convergence. The following constraints were used for fossil time calibrations: (1) Bovinae and Caprinae divergence time (18-22 Ma); (2) Ruminantia and Suina divergence time (48.3-53.5 Ma); (3) Euarchontoglires and Laurasiatheria divergence time (95.3-113 Ma); (4) Euarchontoglires and Rodentia divergence time (85-94 Ma); and (5) Cervus and Elaphurus divergence time (< 3 Ma).

#### Gene family expansions and contractions

The CAFE program (v3.1) [43] was used to analyze gene family expansions and contractions. The program uses a birth and death process to model gene gain and loss across a user-specified phylogenetic tree. The numbers of sika deer genes relative to the number of inferred ancestor genes and expanding and contracting gene families were obtained. According to the GO and KEGG pathway results of the functional annotation, the hypergeometric distribution was used for enrichment analysis, and the BH (Benjamini and Hochberg) algorithm was used for *P*-value correction. A *P*-value less than 0.05 after correction was considered a significant enrichment result.

We investigated several *UGT* genes in each category for the 19 species. The annotated *UGT* genes of human and sika deer were used to predict the unannotated *UGT* genes in the other 17 species with the program GeneWise [40]. MUSCLE software was used for the multiple sequence alignment of all these *UGT* gene protein sequences, whereby a phylogenetic *UGT* gene tree was constructed using RAxML [41].

#### Synteny analysis

A collinearity analysis between sika deer and red deer was conducted using the MUMmer package (v3.23). Furthermore, to identify the synteny block among sika deer, red deer, cattle and goats, we used MCscan (python version) [44] to search for and visualize intragenomic syntenic regions. A homologous synteny block map between sika deer and cattle was plotted with Circos.

#### Demographic history reconstruction

We inferred the demographic histories of sika deer using the Pairwise Sequentially Markovian Coalescent (PSMC) model for diploid genome sequences. The whole-genome diploid consensus sequence for PSMC input was generated by mapping short reads to the sika deer genome with BWA (v0.7.10-r943-dirty) and SAMTools. Program ‘fq2psmcfa’ transforms the consensus sequence into a fasta-like format. The parameters for ‘psmc’ were set as follows: -N25 -t15 -r5 -p “4+25*2+4+6”. The generation times (g) of sika deer, cattle, and buffalo were 5 and 6 years, respectively. The mutation rate for all species was 2.2e-9 per site per year.

#### Positive selection genes

For the single-copy orthologous genes of 19 species, multiple sequence alignment was carried out using MUSCLE (v3.8.31). Regions of uncertain alignment were removed by Gblocks 0.91b [45]. We used branch-site models and likelihood ratio tests (LRTs) in the CODEML of PAML (v4.8a) [42] to detect positive selection genes (PSGs) in the sika deer genome. *P*-values were computed using the χ^2^ statistic and corrected for multiple testing by the false discovery rate (FDR) method (*Padj* < 0.05). All the PSGs were mapped to KEGG pathways and assigned GO terms. GO and KEGG enrichment analyses were then applied to detect the significantly enriched biological processes and signaling pathways of positively selected genes (*Padj* < 0.05).

#### Transcriptome analysis

We performed RNA sequencing of 15 tissues (hypothalamus, liver, muscle, spleen, kidney, testis, pituitary, cecum, duodenum, ileum, jejunum, rumen, abomasum, reticulum and omasum) for each of the 12 sika deer from the feeding trials to determine variations in gene expression levels after treatment. To compare the response to different tannin levels between cattle and sika deer, we conducted RNA-seq and transcriptome analyses of 8 tissues (hypothalamus, liver, kidney, rumen, jejunum, pituitary, reticulum and spleen) from two groups of 6 individuals with a diet containing 0% or 10% gallotannic acid (GA). Total RNA from 226 feeding experiment samples was extracted and used for library construction and sequencing. All libraries were sequenced using an Illumina HiSeq platform.

The transcriptome data of each sample were mapped to the sika deer and cattle genomes using HISAT2 (v2.0.5), and gene expression was calculated in each sample using StringTie (v1.3.0). The R language package DESeq2 was used to homogenize the expression and calculate the differential expression between each pair of samples, in which genes with Padj < 0.05 were considered differentially expressed genes. For the DEGs, the hypergeometric distribution and BH (Benjamin and Hochberg) algorithm were used in the GO and KEGG enrichment analysis and *P*-value correction, respectively. A Q value < 0.05 was considered significantly enriched in the GO and KEGG pathways.

## Authors’ contributions

F.Y., X.X., C.L., and J.R. conceived of the project and designed the research; P.H. drafted the manuscript with input from all authors; C.A., T.W., Y.L., H.T.L., Q.Q. and Q. L. revised the manuscript; C.A., T.W., Y.L. and H.T.L. performed the majority of the analysis, with contributions from H.M. L, R.Z., H.W. and L.W.; Y.C. and S.Z. prepared the library and performed the sequencing; S.W. and A.L. performed the genome assembly with help from W.Z.; T.W. obtained the Hi-C data; X.W. performed the genome annotation analysis; C.A. conducted the positive selection and repeat annotation analysis; X.S. and C.A. performed the gene family analysis; C.A. and T.W. performed the genome collinearity analysis; T.W. performed the reverse transcription analysis; G.W., H.T.L. and J.Z. conducted the feeding trials and prepared the samples for transcriptome sequencing with help from H.W., R.Z., X.S., S.S., Z.Y., T.Y., Y.D., Y.J., L.S., P.Z., H.F., J.X. and X.C.; C.A., M.R., S.C., X.W., W.Y., M.Y.; T.W., and Y.L. performed the analysis of transcriptome data; J.Y. and Y.T. provided the geographic distribution data for Mongolian oak; Y.L. conducted the analysis of the geographic distributions of Mongolian oak and sika deer; C.A., Y.L., T.W. and H.T.L. performed the charting and graphing; and all authors read and approved the final manuscript.

## Acknowledgements

This work was supported by the National Key R&D Program of China (2018YFD0502204), the Agricultural Science and Technology Innovation Program of China (CAAS-ASTIP-2019-ISAPS), the Special Animal Genetic Resources Platform of NSTIC (TZDWZYK2019) and the Sika deer Genome Project of China (20140309016YY).

## Availability of data and material

The whole-genome sequence data reported in this paper have been deposited in the Genome Warehouse in the National Genomics Data Center, Beijing Institute of Genomics (China National Center for Bioinformation), Chinese Academy of Sciences, under accession number GWHANOY00000000, which is publicly accessible at https://bigd.big.ac.cn/gwh. The raw sequence data have been deposited in the Genome Sequence Archive in the National Genomics Data Center under accession numbers CRA001393, CRA002054 and CRA002056, which are publicly accessible at https://bigd.big.ac.cn/gsa.

All procedures concerning animals were performed in accordance with the guidelines for the care and use of experimental animals established by the Ministry of Agriculture of China, and all protocols were approved by the Institutional Animal Care and Use Committee of Institute of Special Economic Animal and Plant Sciences, Chinese Academy of Agricultural Sciences, Changchun, China.

## Supplementary material

**Figure S1 Karyotype of the sequenced female sika deer.** The karyotype analysis shows that the sika deer chromosome number is 2n=66

**Figure S2 Distribution of the 25-mer frequency in the sika deer genome.** The genome size of sika deer is 2.6 Gb based on Kmer analysis with Kmer=25

**Figure S3 Assembly strategy of the sika deer genome.** PacBio long reads were *de novo* assembled with wtdbg2. The chromosome-scale scaffolds were generated by using Hi-C data after genomic error correction. A BioNano optical map and proximal species (red deer) genome were used to check the assembly accuracy

**Figure S4 Genome synteny analysis between sika deer and red deer.** The x-axis represents red deer chromosomes, and the y-axis represents sika deer chromosomes. These two assemblies show significant genomic synteny

**Figure S5 Hi-C interaction heatmap for each chromosome of the sika deer genome**

**Figure S6 Gene syntenic blocks between the sika deer genome and the three ruminant genomes.** The representative chromosome fission/separation fragment is indicated in purple, turquoise and cyan. Gray wedges in the background highlight conserved syntenic blocks with more than 10 gene pairs

**Figure S7 Distribution of identified transposable elements among different mammalian species.** Data anomalies of red deer may be due to the poor quality of the genome

**Figure S8 Circos plot of the chromosomal features of sika deer.** The external green circle represents the chromosomes of sika deer. The circles and links inside the chromosomes from outside to inside represent the distribution of genes in the chromosomes (blue); distribution of repeats of the genome (orange); distribution of heterozygosity (green); and segmental duplications (length >10 kb) (red)

**Figure S9 Phylogeny and divergence time of 19 species.** Maximum-likelihood (ML) tree inferred from single-copy orthologous genes by RAxML. The x-axis is the inferred divergence time (M year) based on the phylogenetic tree and fossils

**Figure S10 Gene family expansion and contraction analysis.** The number of expanded gene families is in red, and the number of contracted gene families is in green

**Figure S11 Phylogenetic tree of all *UGT* genes.** Phylogeny structured by RAxML based on the multiple sequence alignment of all *UGT* genes. These *UGTs* were divided into seven groups. The star represents significantly differentially expressed genes

**Figure S12 Expression heatmap of differentially expressed genes (DEGs) among different treatments**

**Figure S13 Expression of *UGT* genes in 8 tissues of cattle.** *UGT* genes were highly expressed in the liver, kidney and jejunum

**Figure S14 *CYP* gene expression patterns in sika deer.** Five differentially expressed *CYP* genes were upregulated in the liver tissue with increasing tannin intake

**Figure S15 Potential metabolism of drugs and exogenous substances, such as tannins, in the mammalian body.** Oak leaves are rich in hydrolysable tannins. Proline-rich salivary proteins (PRPs) found in the mouth can precipitate gallotannic acid (GA) and play a role in the defense against GA. However, PRPs are not found in all the published genomes of cattle, sheep and our Mhl_v1.0. In the rumen, GA is hydrolyzed into gallic acid and ellagic acid, which are degraded by rumen microbes into simple phenolic compounds. Some of these compounds can be metabolized by the P450 enzyme and excreted from the body. Glucuronyltransferase (GT), sulfatyltransferase (SULT), glutathione S-transferase (GST) and other enzymes produced by the liver can catalyze the conversion of undigested phenolic compounds into glucuronates, sulfates and other water-soluble compounds that can be excreted through the urine. Our results show that only the expression of *UGTs* increased with the tannin content in the liver

**Figure S16 Comparison of the liver, kidney and heart in sika deer, cattle and sheep after a tannin feeding experiment.** The three tissues showed no difference between the treatment group and the control group in sika deer. However, lesions (white arrow) occurred in the three tissues of cattle and sheep. These results demonstrated different tannin tolerances among the 3 species

**Figure S17 Distribution of the insertion segment of Illumina paired-end data**. Illumina sequencing data were generated with four different insert fragment sizes (200, 300, 400, and 600 bp)

**Table S1 Estimation of the sika deer genome size using K-mer analysis**

**Table S2 Summary of the genome sequencing of sika deer**

**Table S3 Summary of the sika deer genome assembly**

**Table S4 Summary of the Hi-C assembly of chromosome length in sika deer**

**Table S5 Summary of the Cervidae genome assembly**

**Table S6 Assessment of the completeness and accuracy of the sika deer genome Table S7 Summary of the repeat content in the sika deer genome**

**Table S8 Comparison of the identified transposable elements among different mammalian species**

**Table S9 Functional annotation of sika deer genes**

**Table S10 Summary of predicted protein-coding genes and gene characteristics**

**Table S11 BUSCO of annotation and assembly**

**Table S12 Statistics for the gene families**

**Table S13 Positively selected genes (PSGs) identified in sika deer**

**Table S14 Functionally enriched KEGG pathway categories of sika deer expanded genes**

**Table S15 Functionally enriched GO categories of sika deer expanded genes Table S16 Functionally enriched KEGG pathway categories of sika deer contracted genes**

**Table S17 Functionally enriched GO categories of sika deer contracted genes**

**Table S18 Numbers of annotated *UGT* genes in 19 species**

**Table S19 Design of the feeding experiment**

