## Supplementary material for "The First High-Quality Reference Genome of Sika Deer Provides Insights for High-Tannin Adaptation": Figure S1

**Supplemental figures**

**Figure S1.** Karyotype of the female sika deer we sequenced 2

**Figure S2.** Distribution of the 25-mer frequency in the sika deer genome 3

**Figure S3.** Assembly strategy of the sika deer genome 4

**Figure S4.** Genome synteny analysis between sika deer and red deer 5

**Figure S5.** Hi-C interaction heatmap for each chromosome of the sika deer genome 6

**Figure S6.** Gene syntenic blocks between the sika deer genome and the three ruminant genomes 7

**Figure S7.** Distribution of identified transposable elements among different mammalian species 8

**Figure S8.** Circos plot of the chromosomal features of sika deer 9

**Figure S9.** Phylogeny and divergence time of 19 species 10

**Figure S10.** Gene family expansion and contraction analysis 11

**Figure S11.** Phylogenetic tree of all *UGT* genes 12

**Figure S12.** Expression heatmap of differentially expressed genes (DEGs) among different treatments 13

**Figure S13.** Expression of *UGT* genes in 8 tissues of cattle 14

**Figure S14.** *CYP* gene expression patterns in sika deer 15

**Figure S15.** Potential metabolism of drugs and exogenous substances, such as tannins, in the mammalian body 16

**Figure S16.** Comparison of liver, kidney and heart in sika deer, cattle and sheep after a tannin feeding experiment 17

**Figure S17.** Distribution of the insertion segment of Illumina paired-end data 18

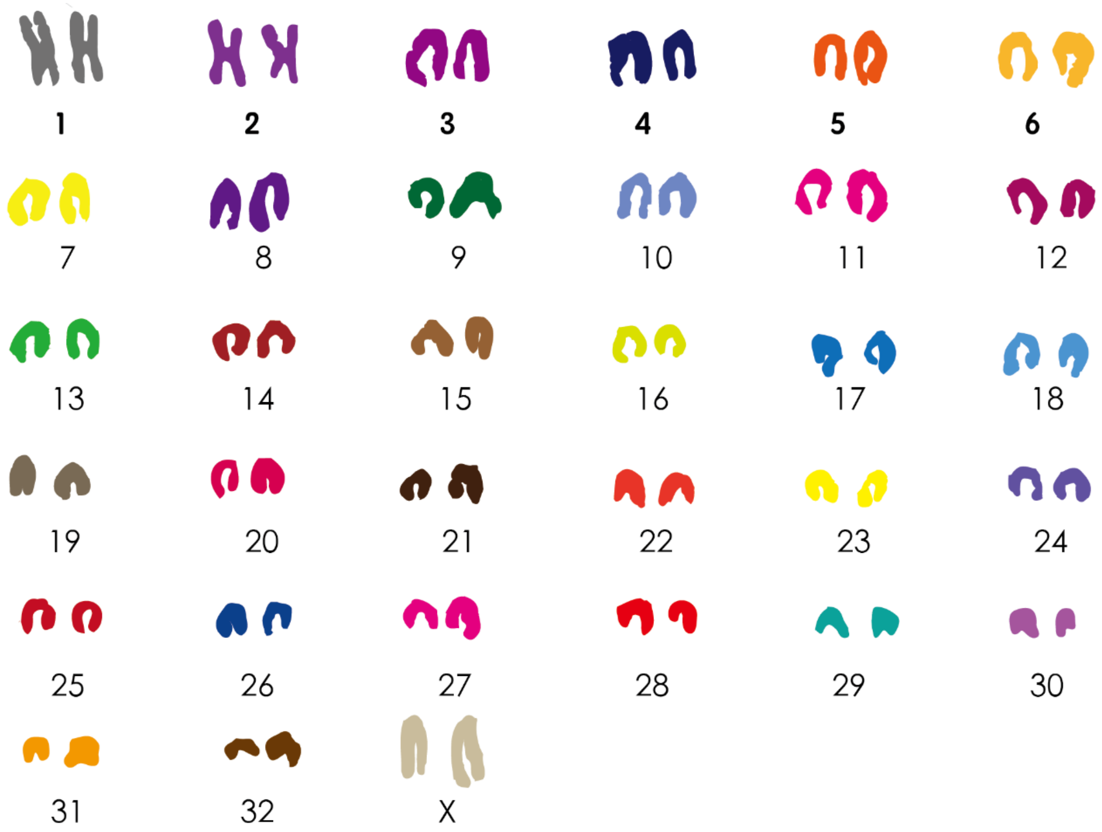


**Figure S1. Karyotype of the sequenced female sika deer.** The karyotype analysis shows that the sika deer chromosome number is 2n=66.

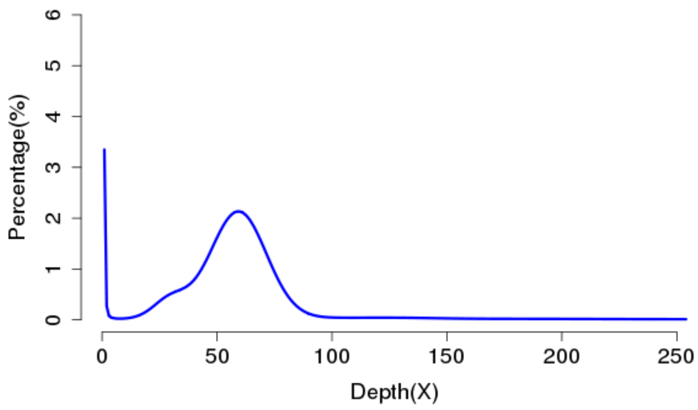


**Figure S2. Distribution of the 25-mer frequency in the sika deer genome.** The genome size of sika deer is 2.6 Gb based on Kmer analysis with Kmer=25.

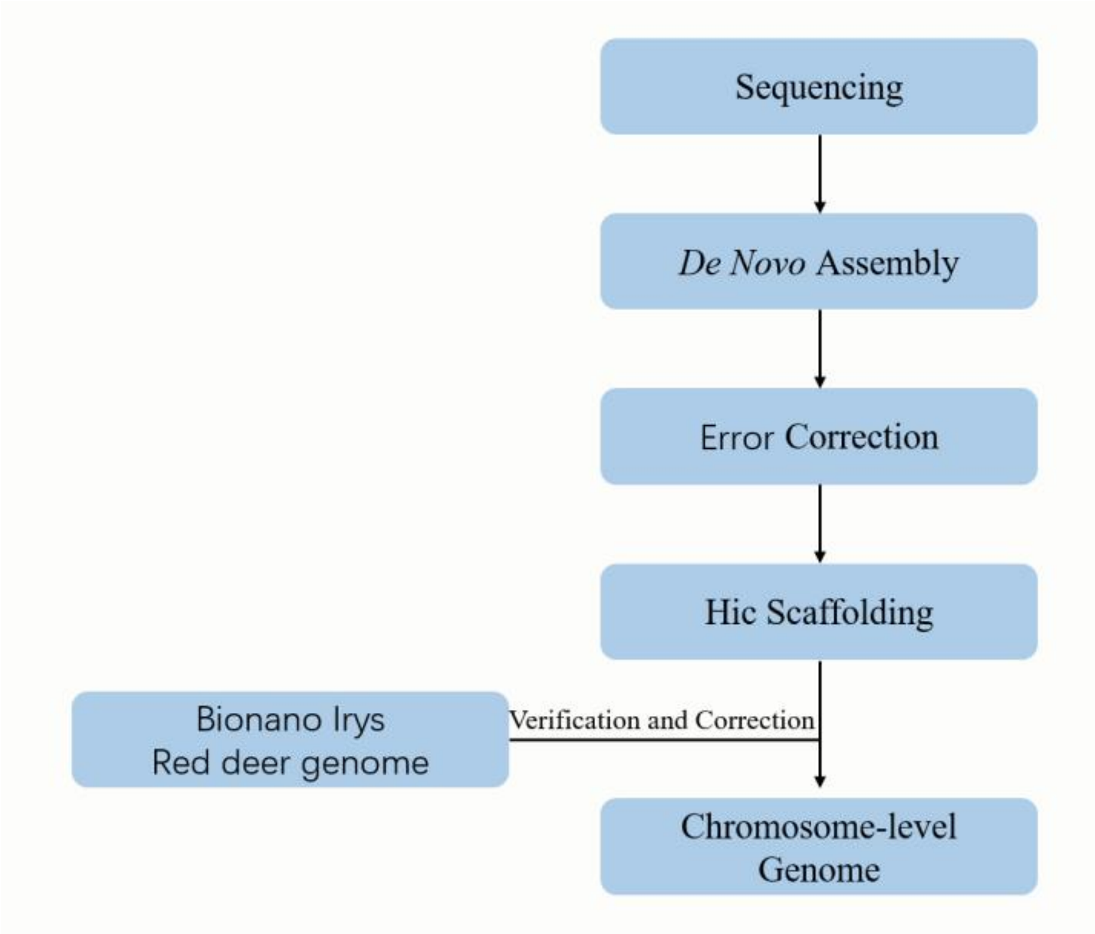


**Figure S3. Assembly strategy of the sika deer genome.** PacBio long reads were *de novo* assembled with wtdbg2. The chromosome-scale scaffolds were generated by using Hi-C data after genomic error correction. A BioNano optical map and proximal species (red deer) genome were used to check the assembly accuracy.

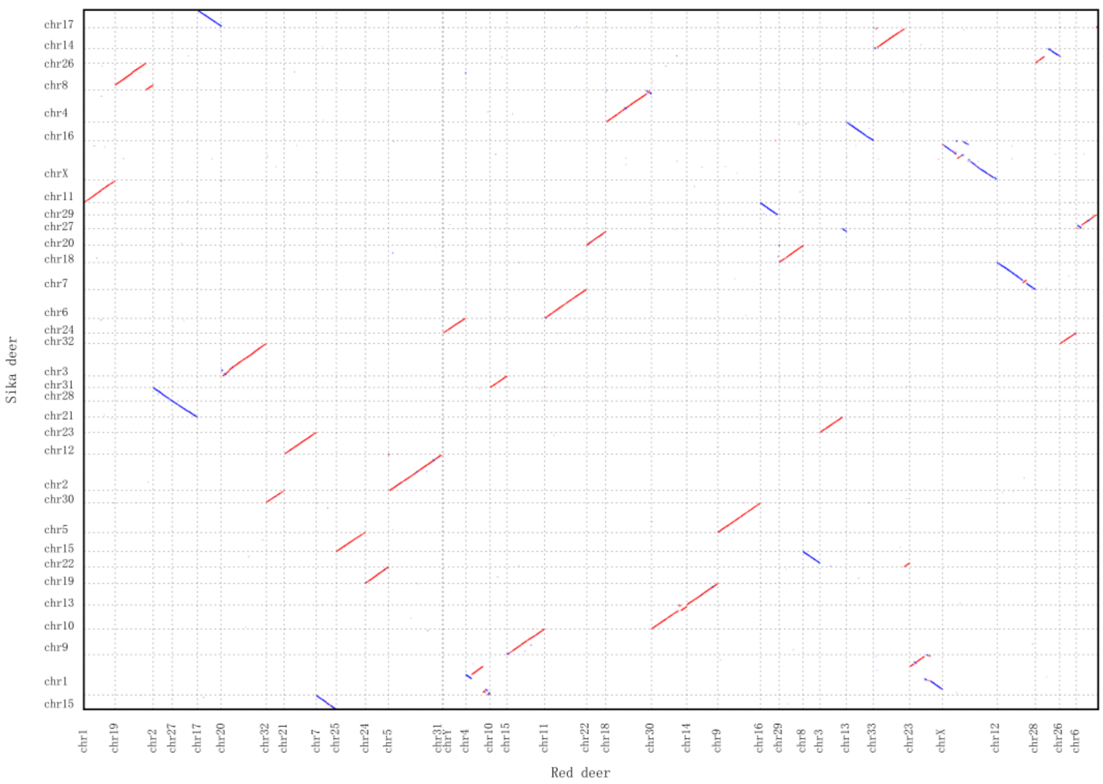


**Figure S4. Genome synteny analysis between sika deer and red deer.** The x-axis represents red deer chromosomes, and the y-axis represents sika deer chromosomes. These two assemblies show significant genomic synteny.

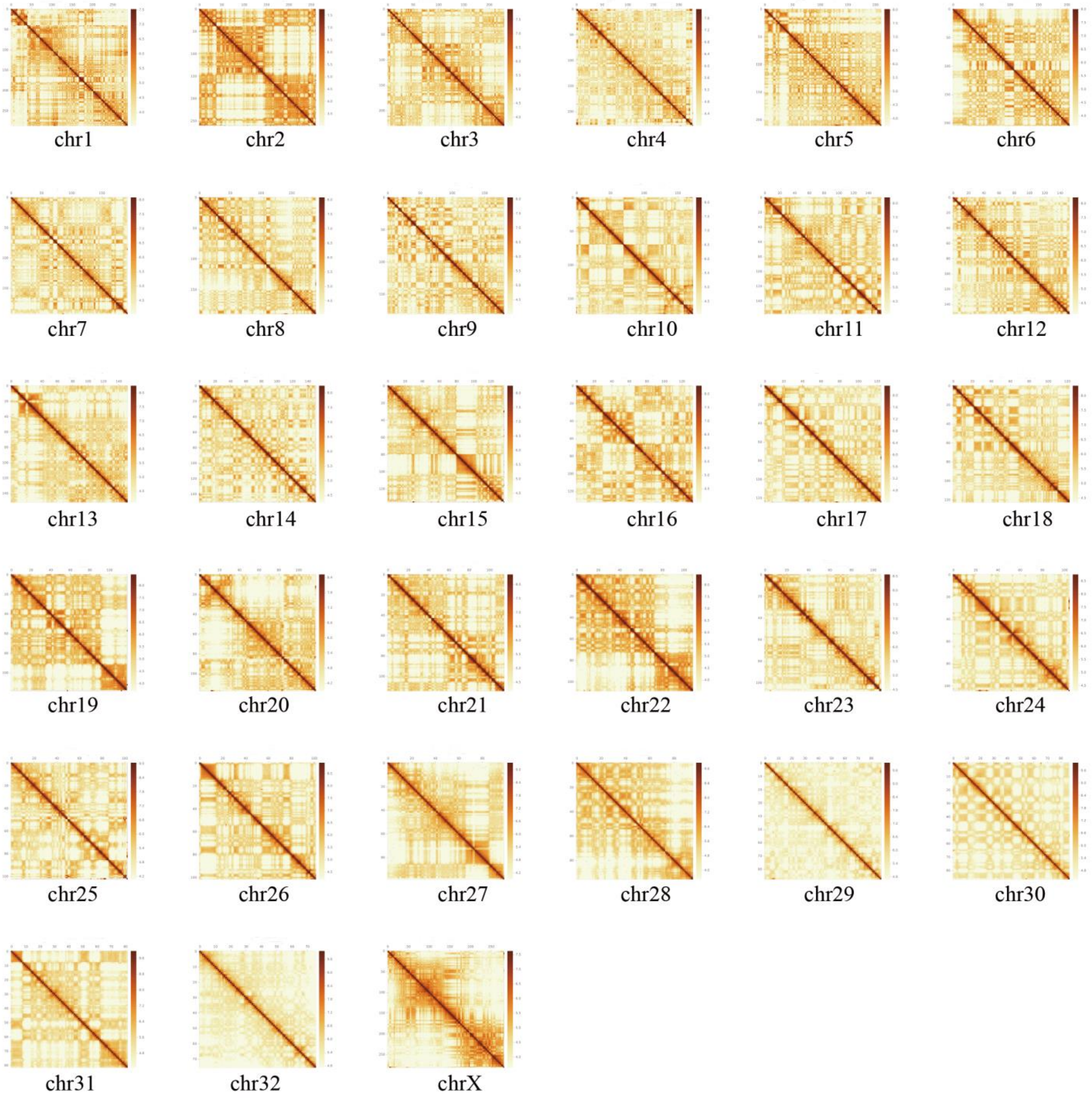


**Figure S5. Hi-C interaction heatmap for each chromosome of the sika deer genome.**

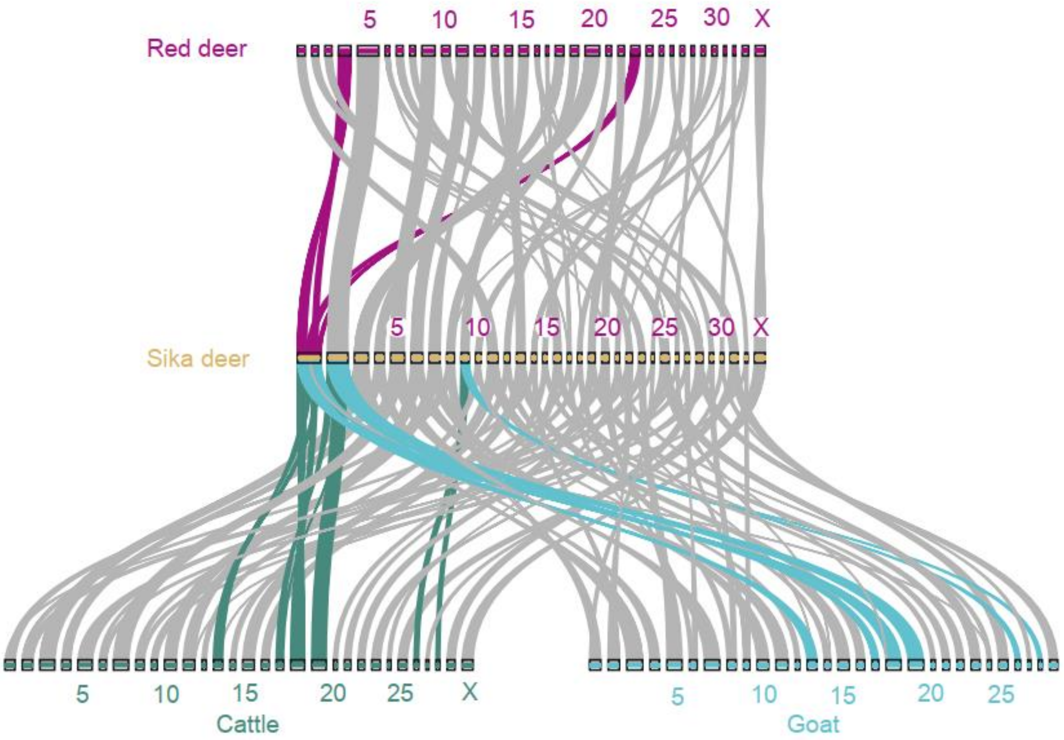


**Figure S6. Gene syntenic blocks between the sika deer genome and the three ruminant genomes.** The representative chromosome fission/separation fragment is indicated in purple, turquoise and cyan. Gray wedges in the background highlight conserved syntenic blocks with more than 10 gene pairs.

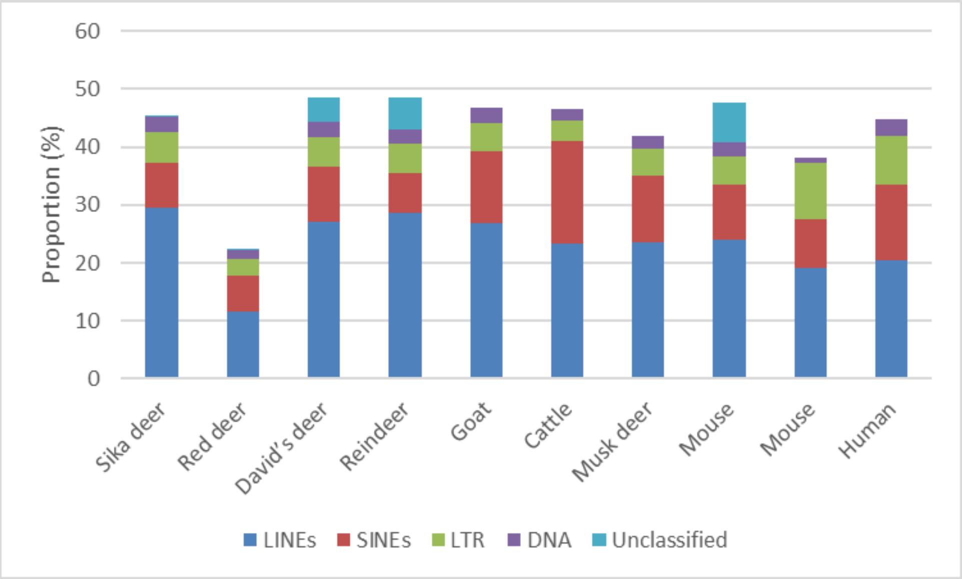


**Figure S7. Distribution of identified transposable elements among different mammalian species.** Data anomalies of red deer may be due to the poor quality of the genome.

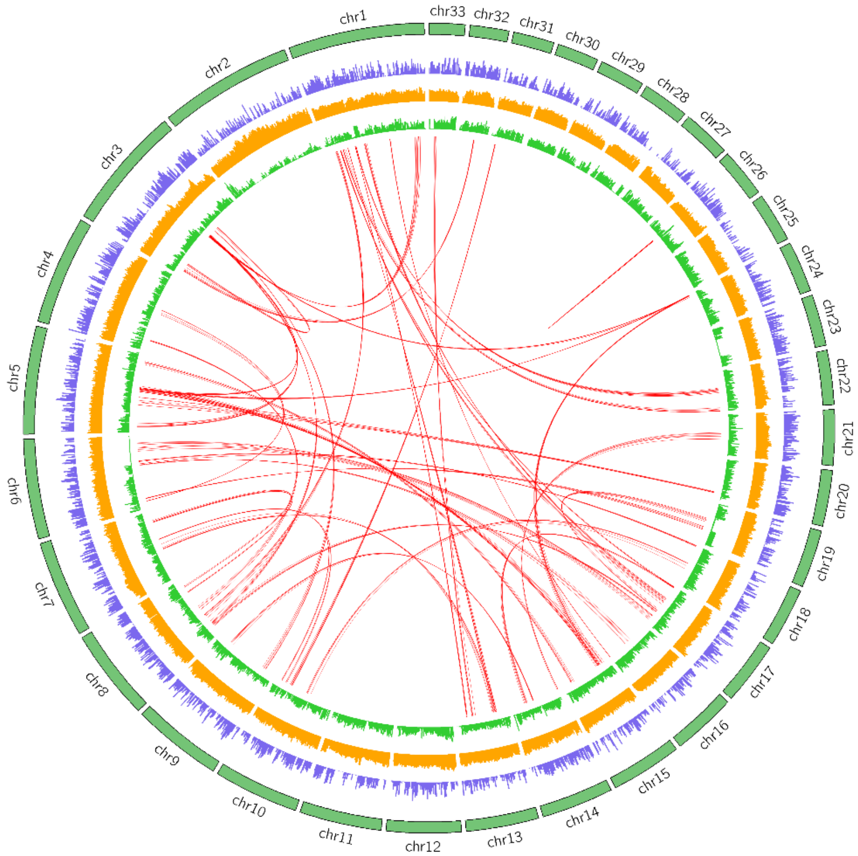


**Figure S8. Circos plot of the chromosomal features of sika deer.** The external green circle represents the chromosomes of sika deer. The circles and links inside the chromosomes from outside to inside represent the distribution of genes in the chromosomes (blue); distribution of repeats of the genome (orange); distribution of heterozygosity (green); and segmental duplications (length >10 kb) (red).

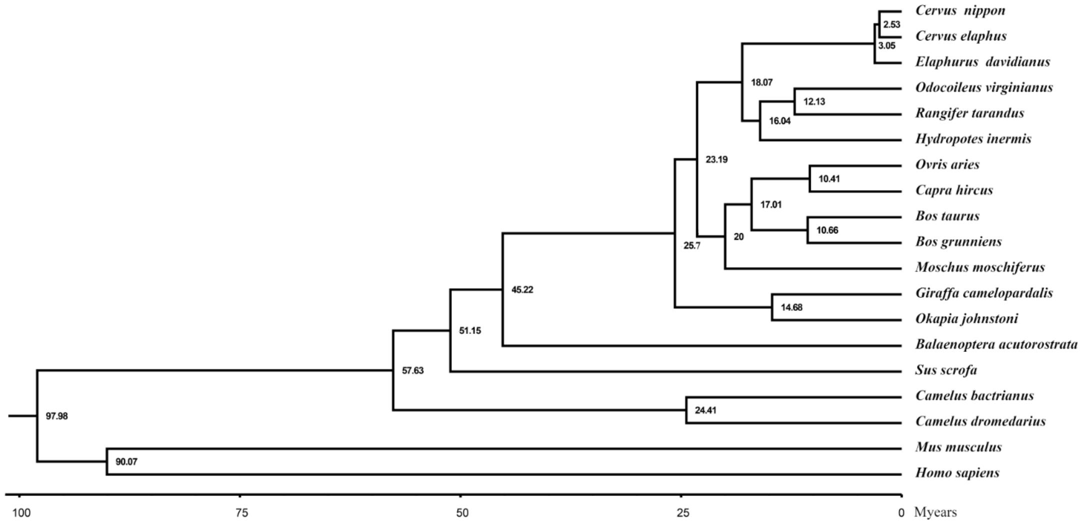


**Figure S9. Phylogeny and divergence time of 19 species.** Maximum-likelihood (ML) tree inferred from single-copy orthologous genes by RAxML. The x-axis is the inferred divergence time (M year) based on the phylogenetic tree and fossils.

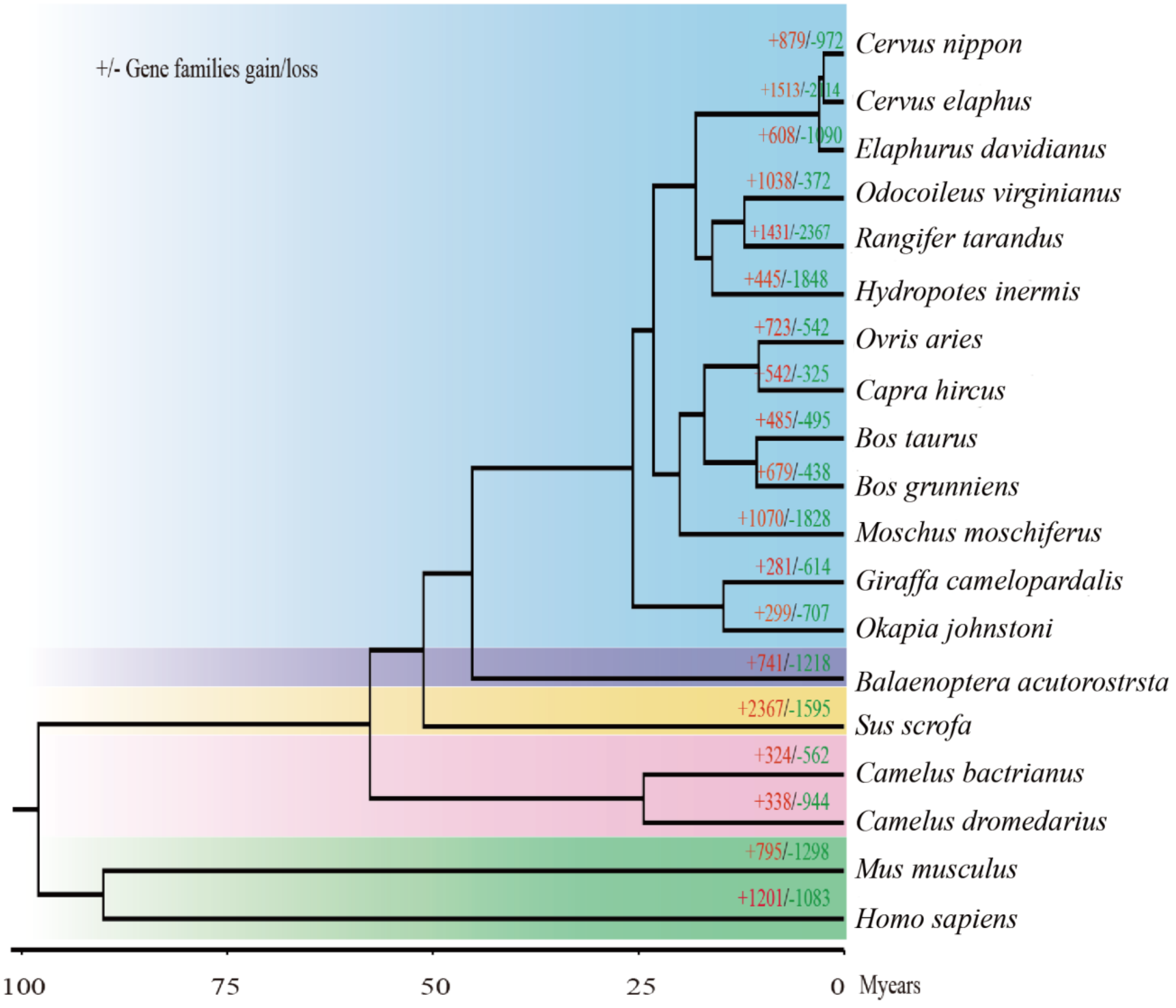


**Figure S10. Gene family expansion and contraction analysis.** The number of expanded gene families is in red, and the number of contracted gene families is in green.

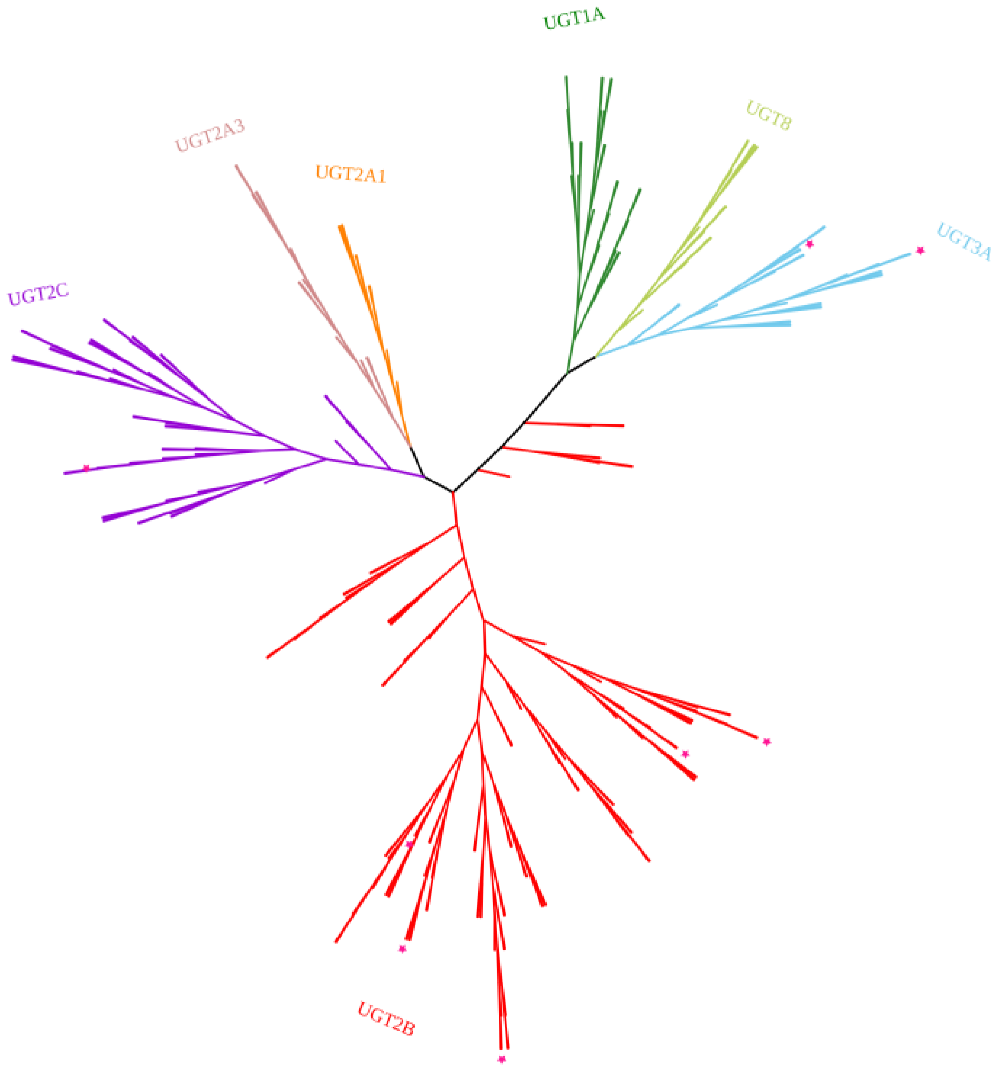


**Figure S11. Phylogenetic tree of all *UGT* genes.** Phylogeny structured by RAxML based on the multiple sequence alignment of all *UGT* genes. These *UGTs* were divided into seven groups. The star represents significantly differentially expressed genes.

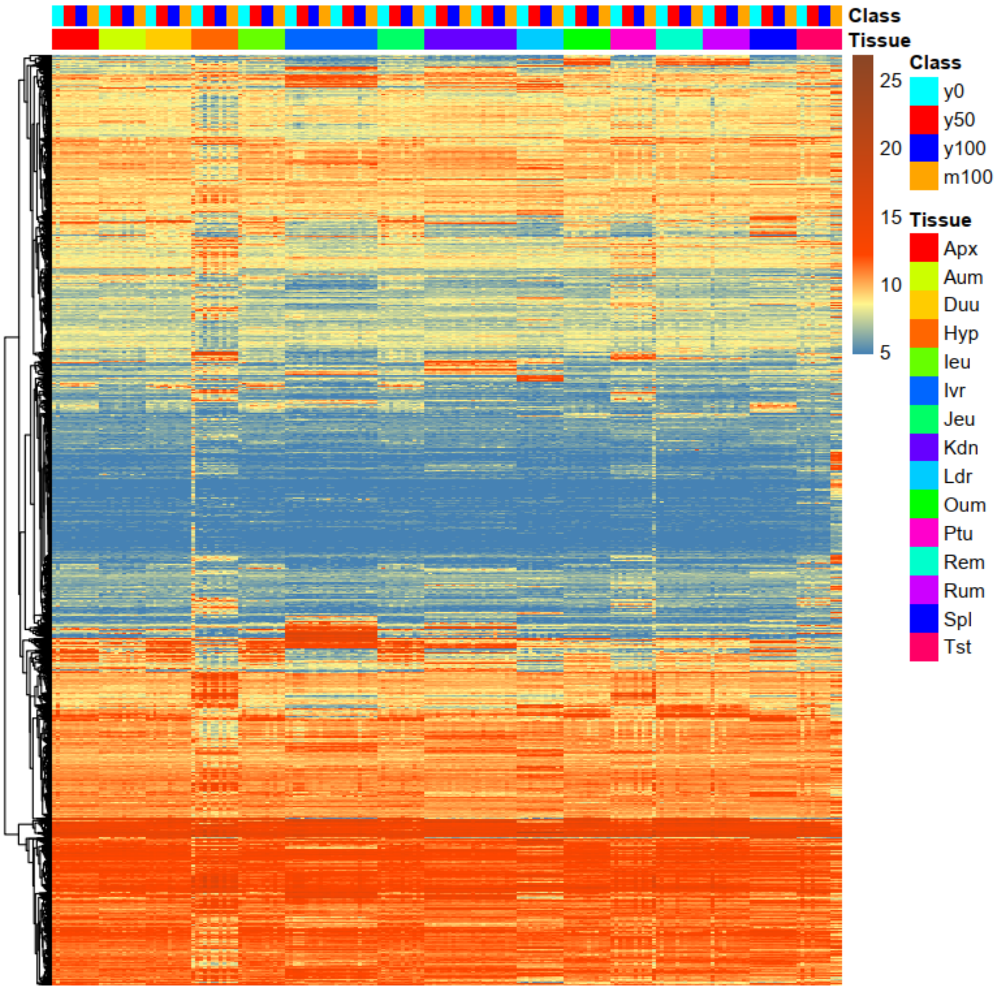


**Figure S12. Expression heatmap of differentially expressed genes (DEGs) among different treatments.**

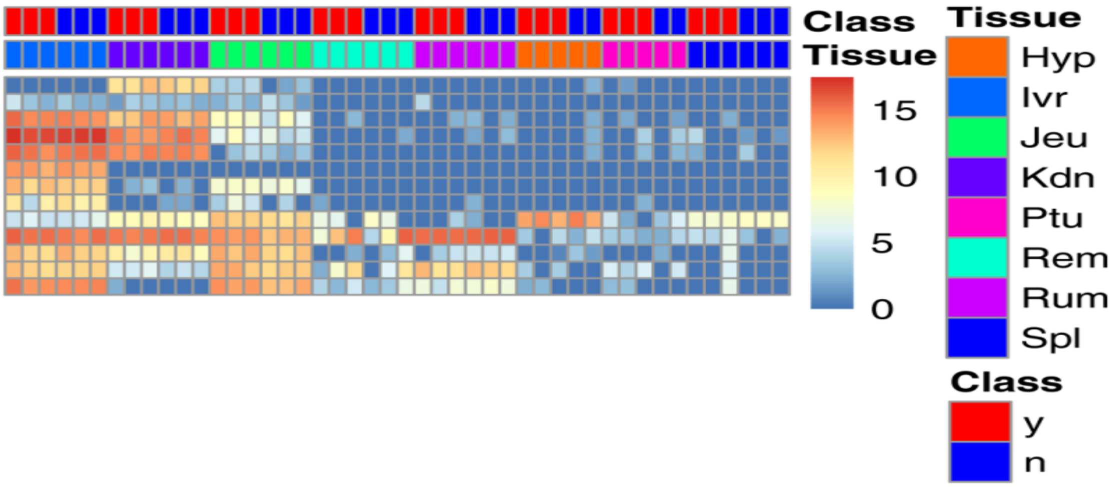


**Figure S13. Expression of *UGT* genes in 8 tissues of cattle.** *UGT* genes were highly expressed in the liver, kidney and jejunum.

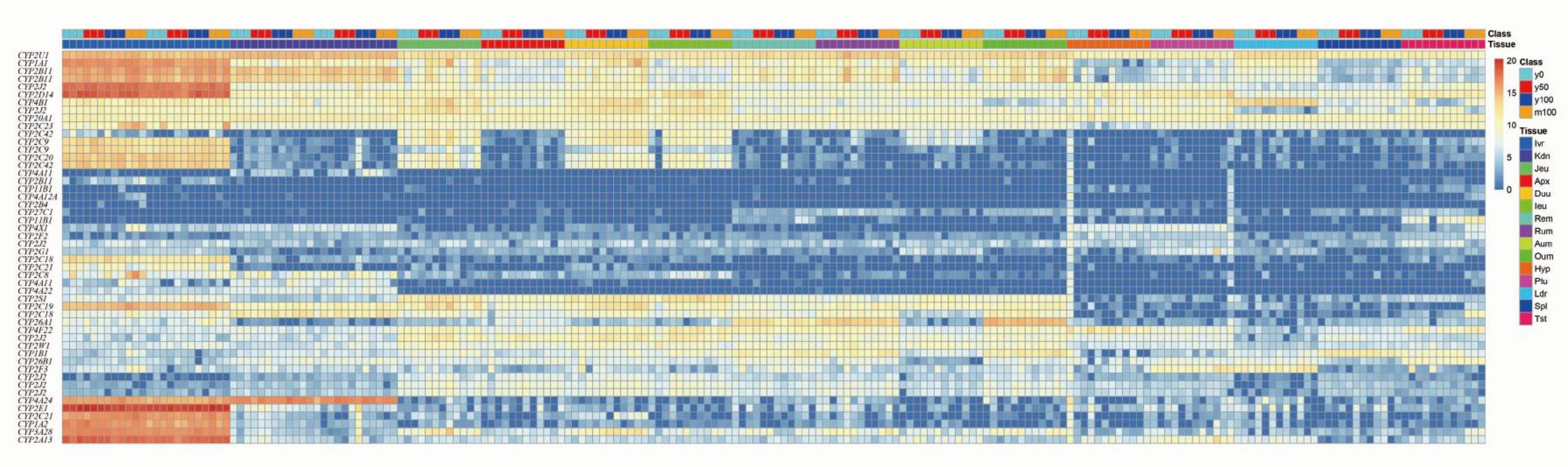


**Figure S14. *CYP* gene expression patterns in sika deer.** Five differentially expressed *CYP* genes were upregulated in the liver tissue with increasing tannin intake.

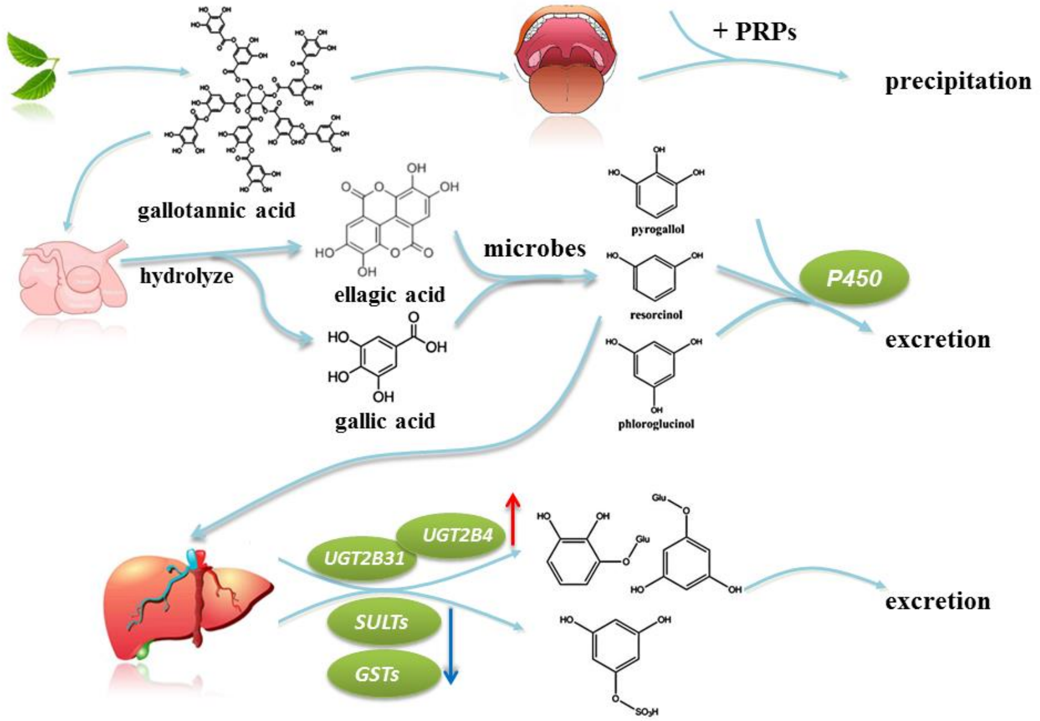


**Figure S15. Potential metabolism of drugs and exogenous substances, such as tannins, in the mammalian body.** Oak leaves are rich in hydrolysable tannins. Proline-rich salivary proteins (PRPs) found in the mouth can precipitate gallotannic acid (GA) and play a role in the defense against GA. However, PRPs are not found in all the published genomes of cattle, sheep and our Mhl_v1.0. In the rumen, GA is hydrolyzed into gallic acid and ellagic acid, which are degraded by rumen microbes into simple phenolic compounds. Some of these compounds can be metabolized by the P450 enzyme and excreted from the body. Glucuronyltransferase (GT), sulfatyltransferase (SULT), glutathione S-transferase (GST) and other enzymes produced by the liver can catalyze the conversion of undigested phenolic compounds into glucuronates, sulfates and other water-soluble compounds that can be excreted through the urine. Our results show that only the expression of *UGTs* increased with the tannin content in the liver.

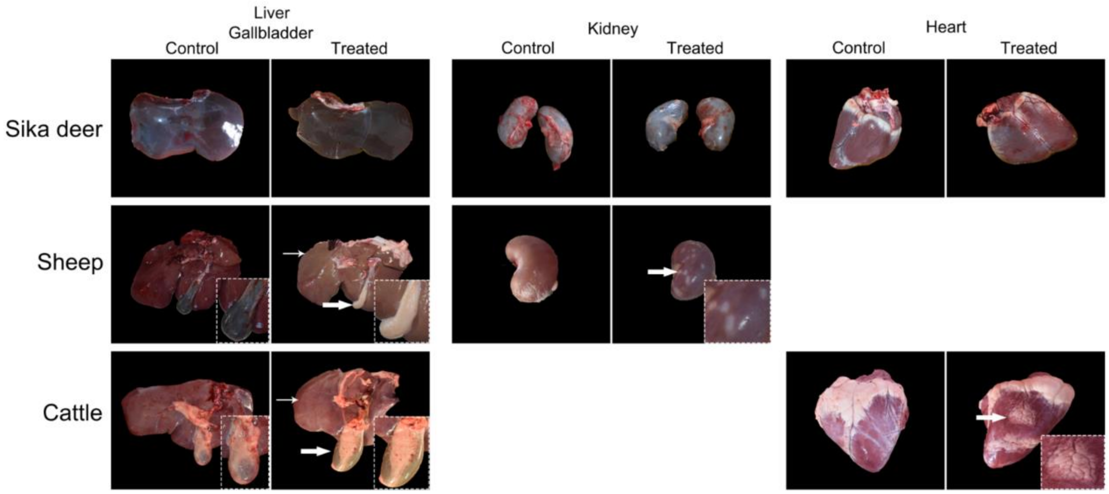


**Figure S16. Comparison of the liver, kidney and heart in sika deer, cattle and sheep after a tannin feeding experiment.** The three tissues showed no difference between the treatment group and the control group in sika deer. However, lesions (white arrow) occurred in the three tissues of cattle and sheep. These results demonstrated different tannin tolerances among the 3 species.

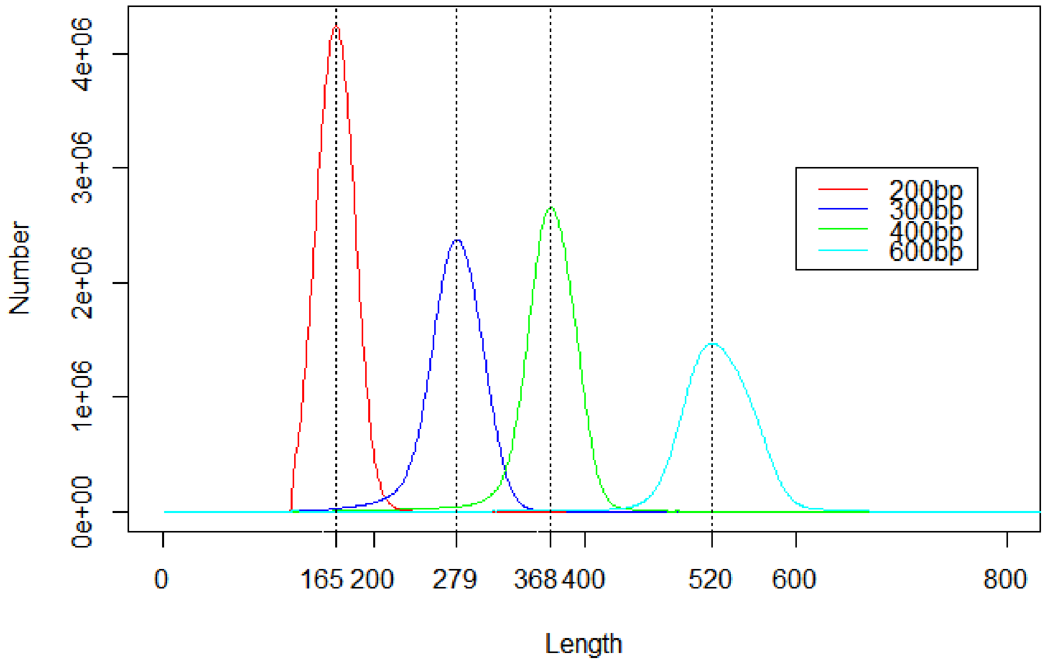


**Figure S17. Distribution of the insertion segment of Illumina paired-end data**. Illumina sequencing data were generated with four different insert fragment sizes (200, 300, 400, and 600 bp).
