## Supplementary material for "The First High-Quality Reference Genome of Sika Deer Provides Insights for High-Tannin Adaptation": Tables S1

**Supplemental tables**

**Table S1.** Estimation of the sika deer genome size using K-mer analysis 2

**Table S2.** Summary of the genome sequencing of sika deer 3

**Table S3.** Summary of the sika deer genome assembly 4

**Table S4.** Summary of the Hi-C assembly of chromosome length in sika deer 5

**Table S5.** Summary of the Cervidae genome assembly 6

**Table S6.** Assessment of the completeness and accuracy of the sika deer genome 7

**Table S7.** Summary of the repeat content in the sika deer genome 8

**Table S8.** Comparison of the identified transposable elements among different mammal species 9

**Table S9.** Functional annotation of sika deer genes 10

**Table S10.** Summary of the predicted protein-coding genes and gene characteristics 11

**Table S11.** BUSCO of annotation and assembly 12

**Table S12.** Statistics for the gene families 13

**Table S13.** Positively selected genes (PSGs) identified in sika deer 14

**Table S14.** Functionally enriched KEGG pathway categories of sika deer expanded genes 16

**Table S15.** Functionally enriched GO categories of sika deer expanded genes 18

**Table S16.** Functionally enriched KEGG pathway categories of sika deer contracted genes 23

**Table S17.** Functionally enriched GO categories of sika deer contracted genes 25

**Table S18.** Numbers of annotated *UGT* genes in 19 species 28

**Table S19.** Design of the feeding experiment 29

**Table S1.** Estimation of the sika deer genome size using K-mer analysis.

| Species | Kmer length | Read Length | Error Ratio (%) | Kmer number (M) | Uniq Kmer Number (M) | Repeat Ratio (%) | First Peak | Genome Size (Mb) | Data Size (Mb) | Kmer Depth | Coverage Depth |
|---|---|---|---|---|---|---|---|---|---|---|---|
| **Sika deer** | 25 | 125 | 0.155 | 154474 | 7764 | 11.9 | 59 | 2614.17 | 244644.28 | 59.09 | 93.58 |

**Table S2.** Summary of the genome sequencing of sika deer.

| platform | Insert size | Read Length (bp) | raw data | | Qualified data | |
|---|---|---|---|---|---|---|
| | | | Total data (G) | Sequence coverage[1] | Total data (G) | Sequence coverage[1] |
| Illumina | 200 bp | 125 bp | 66.8 | 25.7 | 65.1 | 25.0 |
| | 300 bp | 125 bp | 64.6 | 24.8 | 60.5 | 23.3 |
| | 400 bp | 125 bp | 67.2 | 25.8 | 65.3 | 25.1 |
| | 600 bp | 125 bp | 62.9 | 24.2 | 52.0 | 20 |
| | Total | | 261.5 | 100.6 | 242.9 | 93.4 |
| PacBio | | | 150.4 | 57.7 | | |

[1]Kmer analysis estimates the *Cervus nippon* genome size to be 2.6 G.

**Table S3.** Summary of the sika deer genome assembly.

| | PacBio | PacBio + Hi-C |
|---|---|---|
| Total sequence length | 2,500,501,634 | 2,500,646,934 |
| Total sequence number | 2040 | 588 |
| Max sequence length | 93,588,229 | 143,481,735 |
| Average sequence length | 1,225,736 | 4,252,801 |
| N50 | 23,559,432 | 78,786,809 |
| N90 | 2,960,739 | 49,061,596 |

**Table S4.** Summary of the Hi-C assembly of chromosome length in sika deer.

| CHROM | Sika deer chromosome length |
| --- | --- |
| chrX | 140737465 |
| chr1 | 143481735 |
| chr2 | 130042918 |
| chr3 | 114820201 |
| chr4 | 113852600 |
| chr5 | 105445150 |
| chr6 | 102406750 |
| chr7 | 96153163 |
| chr8 | 94958114 |
| chr9 | 90056637 |
| chr10 | 86294865 |
| chr11 | 78786809 |
| chr12 | 76402347 |
| chr13 | 75945954 |
| chr14 | 75108353 |
| chr15 | 67521484 |
| chr16 | 66389940 |
| chr17 | 62175516 |

| | |
|---|---|
| chr18 | 61139481 |
| chr19 | 59074219 |
| chr20 | 58577949 |
| chr21 | 57410528 |
| chr22 | 54302469 |
| chr23 | 53848698 |
| chr24 | 52120049 |
| chr25 | 51225997 |
| chr26 | 50874937 |
| chr27 | 49061596 |
| chr28 | 47929526 |
| chr29 | 43669067 |
| chr30 | 43252544 |
| chr31 | 40873834 |
| chr32 | 37822908 |

**Table S5.** Summary of the Cervidae genome assembly.

| Species | Common name | Genome size (bp) | Ungapped Length (bp) | Anchored Ungapped Length (bp) | Anchored Rate | Contig N50 (bp) | Scaffold N50 (bp) |
|---|---|---|---|---|---|---|---|
| ***Cervus nippon*** | Sika deer | 2,500,646,934 | 2,500,501,634 | 2,481,618,503 | 99.24% | 23,559,432 | 78,786,809 |
| ***Cervus elaphus*** | Red deer | 3,438,623,608 | 1,960,832,178 | 1,928,180,013 | 98.33% | 7,944 | 107,358,006 |
| ***Axis porcinus*** | Hog deer | 2,676,163,295 | 2,635,847,001 | NA | | 172,761 | 20,764,858 |
| ***Hydropotes intermis*** | Chinese water deer | 2,530,176,423 | 2,484,708,650 | NA | | 131,446 | 13,818,975 |
| ***Moschus moschiferus*** | Siberian musk deer | 3,069,608,437 | 2,964,378,037 | NA | | 34,785 | 11,728,851 |
| ***Przewalskium albirostris*** | White-lipped deer | 2,692,225,130 | 2,642,137,551 | NA | | 39,627 | 3,769,372 |
| ***Elaphurus davidianus*** | Milu | 2,584,693,296 | 2,519,836,415 | NA | | 59,950 | 2,844,142 |
| ***Moschus berezovskii*** | Forest musk deer | 2,818,010,997 | 2,669,554,835 | NA | | 57,706 | 2,509,225 |
| ***Rangifer tarandus*** | Reindeer | 2,897,299,934 | 2,896,823,034 | NA | | 77,671 | 1,360,739 |
| ***Muntiacus crinifrons*** | Black muntjac | 2,681,196,170 | 2,675,889,158 | NA | | 8,265 | 1,305,444 |
| ***Muntiacus muntjak*** | Indian muntjac | 2,703,710,594 | 2,607,294,293 | NA | | 23,470 | 1,258,210 |

| | | | | | | | |
|---|---|---|---|---|---|---|---|
| ***Muntiacus reevesi*** | Chinese muntjac | 2,601,888,952 | 2,564,577,671 | NA | | 72,382 | 1,221,377 |
| ***Rangifer tarandus*** | Reindeer | 2,832,785,815 | | NA | | 91,805 | 1,059,113 |
| ***Odocoileus virginianus*** | White-tailed Deer | 2,380,505,687 | 2,359,014,916 | NA | | 122,019 | 850,721 |
| ***Odocoileus hemionus*** | Mule deer | 2,343,701,333 | 2,342,668,817 | NA | | 113,295 | 838,758 |
| ***Tragulus javanicus*** | Lesser mouse-deer | 3,054,942,239 | | NA | | 6,286 | 243,250 |
| ***Moschus berezovskii*** | Forest musk deer | 3,835,628,252 | 3,408,868,026 | NA | | 24,701 | 213,462 |
| ***Moschus chrysogaster*** | alpine musk deer | 4,972,482,505 | 2,241,238,515 | NA | | 3,769 | 100,428 |
| ***Capreolus*** | Western roe deer | 2,785,377,831 | 2,741,850,513 | NA | | 4,167 | 10,458 |
| ***Bos taurus*** | Cattle | 2,715,853,792 | 2,715,825,630 | 2,628,394,923 | 96.78% | 25,896,116 | 103,308,737 |

**Table S6.** Assessment of the completeness and accuracy of the sika deer genome.

| Method | Parameter | Sika deer |
|---|---|---|
| Illumina Reads Mapping and Call SNP[2] | Coverage | 99.29% |
| | Mapping rate | 99.36% |
| | Ratio of heterozygous SNP | 0.38% |
| | Ratio of homozygous SNP[3] | 0.0011% |
| EST mapping | Total number | 2715 |
| | Covered by genome assembly (90%)[4] | 95.95% |
| RNA reads mapping | Total reads number (M) | 1172.55 |
| | Mapping rate | 93.43% |
| BUSCO | Complete BUSCOs (C) | 94.60% |
| | Complete and single-copy BUSCOs (S) | 92.90% |
| | Complete and duplicated BUSCOs (D) | 1.70% |
| | Fragmented BUSCOs (F) | 2.50% |
| | Missing BUSCOs (M) | 2.90% |
| CEGMA | Complete | 97.18% |
| | Partial | 99.19% |

[2]Using bwa, Illumina reads were mapped to the reference genome, and SAMTools was used to call high-quality SNPs.

[3]Predicted error ratio.

[4]Percentage of ESTs mapped to sika deer with coverage > 90%.

**Table S7.** Summary of the repeat content in the sika deer genome.

| | Repbase TEs | | RepeatModeler | | Combined TEs | |
|---|---|---|---|---|---|---|
| | Length (bp) | % | Length (bp) | % | Length (bp) | % |
| DNA | 58615759 | 2.36 | 38091565 | 1.53 | 67122914 | 2.70 |
| LINE | 627094580 | 25.26 | 639177085 | 25.75 | 733702695 | 29.56 |
| LTR | 117747150 | 4.74 | 95014939 | 3.83 | 133587659 | 5.38 |
| SINE | 259074398 | 10.44 | 131258637 | 5.29 | 189393934 | 7.63 |
| Unknown | 918332 | 0.04 | 2436293 | 0.10 | 2650074 | 0.11 |
| Total[5] | 1063450219 | 42.84 | 905978519 | 36.50 | 1126457276 | 45.38 |
| Other[6] | 69041172 | 2.78 | 25717291 | 1.04 | 26376659 | 1.06 |

[5]Total: total interspersed repeats

[6]Other: small RNA, satellites, simple repeats, and low complexity

**Table S8.** Comparison of the identified transposable elements among different mammalian species.

| Species | Common name | LINEs | SINEs | LTR | DNA | Unclassified | Total[5] |
|---|---|---|---|---|---|---|---|
| *Cervus nippon* | Sika deer | 29.56 | 7.63 | 5.38 | 2.70 | 0.11 | 45.38 |
| *Cervus elaphus* | Red deer | 11.64 | 6.18 | 2.86 | 1.45 | 0.02 | 22.15 |
| *Elaphurus davidianus* | Milu | 27.05 | 9.52 | 5.19 | 2.53 | 4.34 | 41.04 |
| *Rangifer tarandus* | Rein deer | 28.63 | 6.79 | 5.26 | 2.25 | 5.63 | 39.18 |
| *Capra hircus* | Goat | 26.85 | 12.35 | 4.90 | 2.61 | NA[7] | 46.71 |
| *Ovis aries* | Sheep | 27.83 | 6.78 | 4.75 | 2.28 | NA | 42.67 |
| *Bubalus bubalis* | Water buffalo | 39.51 | 7.59 | 13.54 | 1.45 | 0.00 | 45.33 |
| *Bos taurus* | Cattle | 23.29 | 17.66 | 3.62 | 1.96 | NA | 46.54 |
| *Moschus berezovskii* | Musk deer | 23.62 | 11.35 | 4.70 | 2.34 | NA | 42.05 |
| *Giraffa camelopardalis* | Giraffe | 24.00 | 9.44 | 4.92 | 2.49 | 6.77 | 39.80 |
| *Mus musculus* | Mouse | 19.20 | 8.22 | 9.87 | 0.88 | 0 | 38.55 |
| *Homo sapiens* | Human | 20.42 | 13.14 | 8.29 | 2.84 | 0 | 44.83 |

[5]Total: total interspersed repeats

[7]NA: repeats not available

**Table S9.** Functional annotation of sika deer genes.

| Category | Gene number | Ratio |
|---|---|---|
| GO | 14161 | 66.0% |
| Interpro | 17889 | 83.4% |
| KEGG | 19208 | 89.6% |
| Swiss-Prot | 18964 | 88.4% |
| TrEMBL | 19246 | 89.7% |
| Total | 19316 | 90.1% |
| Unannotated | 2133 | 9.9% |

**Table S10.** Summary of predicted protein-coding genes and gene characteristics.

| Species | Gene number | Total CDS length (Mb) | Average CDS length | Exon number per gene | Average Exon length |
|---|---|---|---|---|---|
| Sika deer | 21449 | 34.69 | 1617 | 9.29 | 174 |
| Cattle | 19994 | 32.18 | 1609 | 9.64 | 167 |
| Sheep | 20921 | 36.44 | 1597 | 9.91 | 161 |
| Human | 20251 | 28.54 | 1409 | 8.1 | 174 |
| Mouse | 22159 | 32.86 | 1483 | 8.39 | 177 |
| Horse | 20449 | 31.11 | 1521 | 9.22 | 165 |

**Table S11.** BUSCO of annotation and assembly.

| | Complete (C) | Single copy (S) | Duplicated (D) | Fragmented (F) | Missing (M) |
|---|---|---|---|---|---|
| **Annotation** | 3907(95.2%) | 3850(93.8%) | 57(1.4%) | 101(2.5%) | 96(2.3%) |
| **Assembly** | 3879(94.6%) | 3810(92.9%) | 69(1.7%) | 100(2.5%) | 125(2.9%) |

**Table S12.** Statistics for the gene families.

| Species | Common name | Single-copy orthologs | Unique orthologs | Multiple-copy orthologs | Other |
|---|---|---|---|---|---|
| *Homo Sapiens* | Human | 5265 | 22 | 16523 | 678 |
| *Mus musculus* | Mouse | 5390 | 18 | 15743 | 792 |
| *Balaenoptera acutorostrata* | Minke whale | 5303 | 19 | 12275 | 1010 |
| *Camelus bactrianus* | Bactrian camel | 5392 | 18 | 12102 | 1201 |
| *Camelus dromedarius* | Dromedary | 5255 | 0 | 11858 | 1397 |
| *Sus scrofa* | Pig | 4432 | 0 | 16306 | 1266 |
| *Giraffa camelopardalis* | Giraffe | 5368 | 1 | 11472 | 1255 |
| *Okapia johnstoni* | Okapi | 5363 | 0 | 11394 | 1281 |
| *Moschus moschiferus* | Musk deer | 4744 | 27 | 15969 | 1547 |
| *Bos taurus* | Cattle | 5386 | 65 | 13864 | 913 |
| *Bos grunniens* | Yak | 5342 | 51 | 14227 | 857 |
| *Ovris aries* | Sheep | 5276 | 58 | 14402 | 889 |
| *Capra hircus* | Goat | 5416 | 0 | 14382 | 825 |

| | | | | | |
|---|---|---|---|---|---|
| *Odocoileus virginianus* | White-tailed deer | 5251 | 1 | 14745 | 800 |
| *Elaphurus davidianus* | Milu | 5300 | 36 | 12748 | 1133 |
| *Cervus elaphus* | Red deer | 4308 | 1 | 12944 | 1937 |
| *Rangifer tarandus* | Reindeer | 4186 | 2 | 13575 | 2034 |
| *Hydropotes inermis* | Roe deer | 5429 | 0 | 10991 | 1260 |
| *Cervus nippon* | Sika deer | 4918 | 0 | 13486 | 1284 |

**Table S13.** Positively selected genes (PSGs) identified in sika deer.

| symbol | Name | *P*-value |
|---|---|---|
| CDC23 | Cell division cycle protein 23 homolog | 0 |
| BAD | Bcl2-associated agonist of cell death | 0 |
| POLR3H | DNA-directed RNA polymerase III subunit RPC8 | 3E-09 |
| HSPA13 | Heat shock 70 kDa protein 13 | 2.3E-08 |
| DNAJC11 | DnaJ homolog subfamily C member 11 | 0 |
| USHBP1 | Usher syndrome type-1C protein-binding protein 1 | 0 |
| ACAA2 | 3-ketoacyl-CoA thiolase, mitochondrial | 0 |
| TBC1D13 | TBC1 domain family member 13 | 0 |
| JMJD4 | JmjC domain-containing protein 4 | 0 |
| ZW10 | Centromere/kinetochore protein zw10 homolog | 0 |
| EDF1 | Endothelial differentiation-related factor 1 | 0 |
| DDX42 | ATP-dependent RNA helicase DDX42 | 6.1E-08 |
| AGA | N(4)-(beta-N-acetylglucosaminyl)-L-asparaginase | 0 |
| GPN2 | GPN-loop GTPase 2 | 2E-09 |
| SLC25A19 | Mitochondrial thiamine pyrophosphate carrier | 1.45E-03 |
| UFC1 | Ubiquitin-fold modifier-conjugating enzyme 1 | 1.05E-07 |
| PEX12 | Peroxisome assembly protein 12 | 0 |
| ZC3HC1 | Nuclear-interacting partner of ALK | 0 |

| | | |
|---|---|---|
| TGM1 | Protein-glutamine gamma-glutamyltransferase K | 0 |
| Med19 | Mediator of RNA polymerase II transcription subunit 19 | 0 |
| NPAS4 | Neuronal PAS domain-containing protein 4 | 0 |
| TRMT10C | tRNA methyltransferase 10 homolog C | 0 |
| RTF1 | RNA polymerase-associated protein RTF1 homolog | 1.21E-06 |
| RCHY1 | RING finger and CHY zinc finger domain-containing protein 1 | 0 |
| UBLCP1 | Ubiquitin-like domain-containing CTD phosphatase 1 | 0 |
| TOLLIP | Toll-interacting protein | 0 |
| MRPL44 | 39S ribosomal protein L44, mitochondrial | 0 |
| MED1 | Mediator of RNA polymerase II transcription subunit 1 | 0 |
| MORN5 | MORN repeat-containing protein 5 | 0 |
| ZMYND12 | Zinc finger MYND domain-containing protein 12 | 1.51E-03 |
| Pkdcc | Extracellular tyrosine-protein kinase PKDCC | 0 |
| TMF1 | TATA element modulatory factor | 0 |
| PPP1R36 | Protein phosphatase 1 regulatory subunit 36 | 0 |
| PRKCSH | Glucosidase 2 subunit beta | 2.24E-05 |
| MLH3 | DNA mismatch repair protein Mlh3 | 0 |
| - | UPF0449 protein C19orf25 homolog | 5.14E-06 |
| INHA | Inhibin alpha chain | 0 |
| CMBL | Carboxymethylenebutenolidase homolog | 7.85E-04 |

| | | |
|---|---|---|
| TTC22 | Tetratricopeptide repeat protein 22 | 3.44E-05 |
| NKAPD1 | Uncharacterized protein NKAPD1 | 0 |
| DNAJB5 | DnaJ homolog subfamily B member 5 | 0 |
| TBATA | Protein TBATA | 0 |
| RCSD1 | CapZ-interacting protein | 0 |
| CD34 | Hematopoietic progenitor cell antigen CD34 | 2.29E-05 |
| KCNE2 | Potassium voltage-gated channel subfamily E member 2 | 5E-09 |
| ACRBP | Acrosin-binding protein (Fragment) | 0 |
| CD72 | B-cell differentiation antigen CD72 | 9.1E-08 |
| DDN | Dendrin | 0 |
| CCDC155 | Protein KASH5 | 0 |
| Slfnl1 | Schlafen-like protein 1 | 0 |
| TMEM139 | Transmembrane protein 139 | 0 |
| IZUMO3 | Izumo sperm-egg fusion protein 3 | 0 |
| HINFP | Histone H4 transcription factor | 0 |
| DHX30 | Putative ATP-dependent RNA helicase DHX30 | 0 |
| GRAMD1C | GRAM domain-containing protein 1C | 0 |

**Table S14.** Functionally enriched KEGG pathway categories of sika deer expanded genes.

| ID | Categories | *P*-value | *Padj* |
|---|---|---|---|
| ko04740 | Olfactory transduction | 0 | 0 |
| ko03010 | Ribosome | 1.13E-32 | 1.31E-30 |
| ko03011 | Ribosome | 1.13E-32 | 1.31E-30 |
| ko04147 | Exosome | 6.18E-13 | 5.39E-11 |
| ko03000 | Transcription factors | 8.48E-13 | 5.92E-11 |
| ko05034 | Alcoholism | 1.07E-12 | 6.24E-11 |
| ko04812 | Cytoskeleton proteins | 2.41E-12 | 1.20E-10 |
| ko03036 | Chromosome and associated proteins | 5.62E-12 | 2.45E-10 |
| ko04030 | G protein-coupled receptors | 6.66E-12 | 2.58E-10 |
| ko05322 | Systemic lupus erythematosus | 4.86E-11 | 1.70E-09 |
| ko04031 | GTP-binding proteins | 1.23E-09 | 3.89E-08 |
| ko00140 | Steroid hormone biosynthesis | 5.21E-08 | 1.51E-06 |
| ko00310 | Lysine degradation | 6.00E-08 | 1.61E-06 |
| ko04390 | Hippo signaling pathway | 1.05E-07 | 2.61E-06 |
| ko04612 | Antigen processing and presentation | 1.54E-07 | 3.59E-06 |
| ko04626 | Plant-pathogen interaction | 1.78E-07 | 3.87E-06 |
| ko00830 | Retinol metabolism | 2.10E-07 | 4.32E-06 |
| ko05150 | Staphylococcus aureus infection | 6.99E-07 | 1.35E-05 |

| | | | |
|---|---|---|---|
| ko00982 | Drug metabolism - cytochrome P450 | 4.17E-06 | 7.65E-05 |
| ko00053 | Ascorbate and aldarate metabolism | 5.24E-06 | 9.14E-05 |
| ko05310 | Asthma | 6.50E-06 | 1.08E-04 |
| ko04145 | Phagosome | 9.35E-06 | 1.48E-04 |
| ko00040 | Pentose and glucuronate interconversions | 1.35E-05 | 2.05E-04 |
| ko01009 | Protein phosphatase and associated proteins | 1.99E-05 | 2.90E-04 |
| ko05130 | Pathogenic Escherichia coli infection | 4.98E-05 | 6.96E-04 |
| ko03019 | Messenger RNA Biogenesis | 5.43E-05 | 7.29E-04 |
| ko00860 | Porphyrin and chlorophyll metabolism | 1.15E-04 | 1.48E-03 |
| ko03051 | Proteasome | 1.18E-04 | 1.48E-03 |
| ko04350 | TGF-beta signaling pathway | 1.29E-04 | 1.56E-03 |
| ko04672 | Intestinal immune network for IgA production | 1.75E-04 | 2.04E-03 |
| ko05031 | Amphetamine addiction | 2.03E-04 | 2.29E-03 |
| ko04744 | Phototransduction | 2.26E-04 | 2.46E-03 |
| ko04540 | Gap junction | 2.39E-04 | 2.53E-03 |
| ko05332 | Graft-versus-host disease | 2.68E-04 | 2.76E-03 |
| ko04110 | Cell cycle | 2.92E-04 | 2.90E-03 |
| ko04391 | Hippo signaling pathway - fly | 3.00E-04 | 2.90E-03 |
| ko05323 | Rheumatoid arthritis | 3.07E-04 | 2.90E-03 |
| ko00980 | Metabolism of xenobiotics by cytochrome P450 | 3.40E-04 | 3.13E-03 |

| | | | |
|---|---|---|---|
| ko04940 | Type I diabetes mellitus | 3.91E-04 | 3.50E-03 |
| ko05416 | Viral myocarditis | 4.40E-04 | 3.84E-03 |
| ko00534 | Glycosaminoglycan biosynthesis - heparan sulfate/heparin | 5.16E-04 | 4.39E-03 |
| ko00512 | Mucin type O-glycan biosynthesis | 5.98E-04 | 4.86E-03 |
| ko05033 | Nicotine addiction | 5.98E-04 | 4.86E-03 |
| ko04114 | Oocyte meiosis | 7.09E-04 | 5.50E-03 |
| ko05030 | Cocaine addiction | 6.95E-04 | 5.50E-03 |
| ko00983 | Drug metabolism - other enzymes | 9.07E-04 | 6.88E-03 |
| ko01020 | Enzyme-linked receptors | 9.34E-04 | 6.93E-03 |
| ko05012 | Parkinson's disease | 9.77E-04 | 6.96E-03 |
| ko05204 | Chemical carcinogenesis | 9.69E-04 | 6.96E-03 |
| ko05010 | Alzheimer's disease | 1.00E-03 | 7.01E-03 |
| ko00604 | Glycosphingolipid biosynthesis - ganglio series | 1.18E-03 | 8.11E-03 |
| ko00190 | Oxidative phosphorylation | 1.30E-03 | 8.47E-03 |
| ko04218 | Cellular senescence | 1.27E-03 | 8.47E-03 |
| ko05330 | Allograft rejection | 1.31E-03 | 8.47E-03 |
| ko05016 | Huntington's disease | 1.40E-03 | 8.87E-03 |
| ko05166 | HTLV-I infection | 1.45E-03 | 9.04E-03 |
| ko04016 | MAPK signaling pathway - plant | 1.49E-03 | 9.09E-03 |

| | | | |
|---|---|---|---|
| ko05320 | Autoimmune thyroid disease | 2.39E-03 | 1.44E-02 |
| ko04730 | Long-term depression | 2.88E-03 | 1.71E-02 |
| ko00910 | Nitrogen metabolism | 3.06E-03 | 1.78E-02 |
| ko04728 | Dopaminergic synapse | 4.09E-03 | 2.34E-02 |
| ko04015 | Rap1 signaling pathway | 4.38E-03 | 2.46E-02 |
| ko04261 | Adrenergic signaling in cardiomyocytes | 5.24E-03 | 2.90E-02 |
| ko03009 | Ribosome biogenesis | 5.33E-03 | 2.91E-02 |
| ko01003 | Glycosyltransferases | 5.52E-03 | 2.97E-02 |
| ko03041 | Spliceosome | 5.65E-03 | 2.99E-02 |
| ko04516 | Cell adhesion molecules and their ligands | 6.01E-03 | 3.13E-02 |
| ko05321 | Inflammatory bowel disease (IBD) | 6.40E-03 | 3.28E-02 |
| ko04745 | Phototransduction – fly | 9.16E-03 | 4.63E-02 |

**Table S15.** Functionally enriched GO categories of sika deer expanded genes.

| ID | Categories | *P*-value | *Padj* |
|---|---|---|---|
| GO:0003676 | nucleic acid binding | 0 | 0 |
| GO:0004984 | olfactory receptor activity | 0 | 0 |
| GO:0006355 | regulation of transcription, DNA-templated | 0 | 0 |
| GO:0007186 | G-protein coupled receptor signaling pathway | 0 | 0 |
| GO:0046872 | metal ion binding | 0 | 0 |
| GO:0005840 | ribosome | 7.00E-50 | 7.13E-48 |
| GO:0003735 | structural constituent of ribosome | 8.70E-47 | 7.59E-45 |
| GO:0006412 | translation | 1.56E-44 | 1.19E-42 |
| GO:0005882 | intermediate filament | 7.16E-35 | 4.86E-33 |
| GO:0019068 | virion assembly | 1.13E-31 | 6.92E-30 |
| GO:0005198 | structural molecule activity | 9.70E-31 | 5.39E-29 |
| GO:0016887 | ATPase activity | 1.65E-22 | 8.38E-21 |
| GO:0016032 | viral process | 5.34E-22 | 2.51E-20 |
| GO:0045095 | keratin filament | 1.56E-21 | 6.81E-20 |
| GO:0008076 | voltage-gated potassium channel complex | 1.04E-19 | 4.25E-18 |
| GO:0042626 | ATPase activity, coupled to transmembrane movement of substances | 2.73E-19 | 1.04E-17 |
| GO:0019028 | viral capsid | 7.47E-13 | 2.68E-11 |

| | | | |
|---|---|---|---|
| GO:0005249 | voltage-gated potassium channel activity | 9.79E-13 | 3.32E-11 |
| GO:0005622 | intracellular | 2.75E-12 | 8.84E-11 |
| GO:0003700 | transcription factor activity, sequence-specific DNA binding | 3.03E-12 | 9.25E-11 |
| GO:0004930 | G-protein coupled receptor activity | 5.08E-12 | 1.48E-10 |
| GO:0007156 | homophilic cell adhesion via plasma membrane adhesion molecules | 4.89E-11 | 1.36E-09 |
| GO:0000786 | nucleosome | 1.96E-10 | 5.22E-09 |
| GO:0004890 | GABA-A receptor activity | 3.34E-10 | 8.49E-09 |
| GO:0042613 | MHC class II protein complex | 5.78E-10 | 1.41E-08 |
| GO:0016021 | integral component of membrane | 1.45E-09 | 3.41E-08 |
| GO:0043565 | sequence-specific DNA binding | 3.88E-09 | 8.78E-08 |
| GO:0004888 | transmembrane signaling receptor activity | 4.05E-09 | 8.85E-08 |
| GO:0005887 | integral component of plasma membrane | 5.79E-09 | 1.22E-07 |
| GO:0016020 | membrane | 8.43E-09 | 1.72E-07 |
| GO:0016712 | oxidoreductase activity, acting on paired donors, with incorporation or reduction of molecular oxygen, reduced flavin or flavoprotein as one donor, and incorporation of one atom of oxygen | 9.81E-09 | 1.93E-07 |
| GO:0007166 | cell surface receptor signaling pathway | 2.26E-08 | 4.32E-07 |
| GO:0005886 | plasma membrane | 2.57E-08 | 4.77E-07 |

| | | | |
|---|---|---|---|
| GO:0004523 | RNA-DNA hybrid ribonuclease activity | 4.00E-08 | 7.18E-07 |
| GO:0006810 | transport | 4.34E-08 | 7.57E-07 |
| GO:0019904 | protein domain specific binding | 4.69E-08 | 7.96E-07 |
| GO:0051260 | protein homooligomerization | 5.21E-08 | 8.60E-07 |
| GO:0006813 | potassium ion transport | 8.70E-08 | 1.40E-06 |
| GO:0019001 | guanyl nucleotide binding | 1.78E-07 | 2.71E-06 |
| GO:0031683 | G-protein beta/gamma-subunit complex binding | 1.78E-07 | 2.71E-06 |
| GO:0016758 | transferase activity, transferring hexosyl groups | 2.03E-07 | 3.03E-06 |
| GO:0019882 | antigen processing and presentation | 3.74E-07 | 5.43E-06 |
| GO:0004012 | phospholipid-translocating ATPase activity | 6.88E-07 | 9.55E-06 |
| GO:0015914 | phospholipid transport | 6.88E-07 | 9.55E-06 |
| GO:0004983 | neuropeptide Y receptor activity | 1.08E-06 | 1.44E-05 |
| GO:0008021 | synaptic vesicle | 1.08E-06 | 1.44E-05 |
| GO:0046982 | protein heterodimerization activity | 1.16E-06 | 1.51E-05 |
| GO:0005003 | ephrin receptor activity | 2.56E-06 | 3.26E-05 |
| GO:0005525 | GTP binding | 2.65E-06 | 3.31E-05 |
| GO:0007264 | small GTPase-mediated signal transduction | 4.37E-06 | 5.35E-05 |
| GO:0007214 | gamma-aminobutyric acid signaling pathway | 4.99E-06 | 5.98E-05 |
| GO:0048013 | ephrin receptor signaling pathway | 9.58E-06 | 1.13E-04 |
| GO:0005230 | extracellular ligand-gated ion channel activity | 1.30E-05 | 1.50E-04 |

| | | | |
|---|---|---|---|
| GO:0051082 | unfolded protein binding | 1.54E-05 | 1.75E-04 |
| GO:0004180 | carboxypeptidase activity | 2.30E-05 | 2.47E-04 |
| GO:0004459 | L-lactate dehydrogenase activity | 2.30E-05 | 2.47E-04 |
| GO:0005044 | scavenger receptor activity | 2.32E-05 | 2.47E-04 |
| GO:0006811 | ion transport | 2.34E-05 | 2.47E-04 |
| GO:0000166 | nucleotide binding | 2.65E-05 | 2.75E-04 |
| GO:0003924 | GTPase activity | 3.67E-05 | 3.68E-04 |
| GO:0007188 | adenylate cyclase-modulating G-protein coupled receptor signaling pathway | 3.63E-05 | 3.68E-04 |
| GO:0004114 | 3',5'-cyclic-nucleotide phosphodiesterase activity | 3.89E-05 | 3.72E-04 |
| GO:0004143 | diacylglycerol kinase activity | 3.86E-05 | 3.72E-04 |
| GO:0007205 | protein kinase C-activating G-protein coupled receptor signaling pathway | 3.86E-05 | 3.72E-04 |
| GO:0005667 | transcription factor complex | 5.64E-05 | 5.15E-04 |
| GO:0007155 | cell adhesion | 5.65E-05 | 5.15E-04 |
| GO:0050661 | NADP binding | 5.64E-05 | 5.15E-04 |
| GO:0008009 | chemokine activity | 6.60E-05 | 5.93E-04 |
| GO:0005874 | microtubule | 7.30E-05 | 6.37E-04 |
| GO:0007017 | microtubule-based process | 7.30E-05 | 6.37E-04 |
| GO:0007169 | transmembrane receptor protein tyrosine kinase | 7.60E-05 | 6.54E-04 |

| | | | |
|---|---|---|---|
| | signaling pathway | | |
| GO:0003956 | NAD(P)+-protein-arginine ADP-ribosyltransferase activity | 1.06E-04 | 8.39E-04 |
| GO:0004952 | dopamine neurotransmitter receptor activity | 1.06E-04 | 8.39E-04 |
| GO:0004977 | melanocortin receptor activity | 1.06E-04 | 8.39E-04 |
| GO:0005549 | odorant binding | 1.06E-04 | 8.39E-04 |
| GO:0008097 | 5S rRNA binding | 1.06E-04 | 8.39E-04 |
| GO:0016907 | G-protein coupled acetylcholine receptor activity | 1.06E-04 | 8.39E-04 |
| GO:0000785 | chromatin | 1.49E-04 | 1.15E-03 |
| GO:0031492 | nucleosomal DNA binding | 1.49E-04 | 1.15E-03 |
| GO:0004181 | metallocarboxypeptidase activity | 1.60E-04 | 1.22E-03 |
| GO:0019058 | viral life cycle | 2.26E-04 | 1.70E-03 |
| GO:0016705 | oxidoreductase activity, acting on paired donors, with incorporation or reduction of molecular oxygen | 2.39E-04 | 1.78E-03 |
| GO:0008146 | sulfotransferase activity | 2.42E-04 | 1.78E-03 |
| GO:0006935 | chemotaxis | 2.61E-04 | 1.90E-03 |
| GO:0009607 | response to biotic stimulus | 3.26E-04 | 2.34E-03 |
| GO:0005581 | collagen trimer | 4.86E-04 | 3.38E-03 |
| GO:0006913 | nucleocytoplasmic transport | 4.86E-04 | 3.38E-03 |
| GO:0045296 | cadherin binding | 4.86E-04 | 3.38E-03 |

| | | | |
|---|---|---|---|
| GO:0004950 | chemokine receptor activity | 5.08E-04 | 3.49E-03 |
| GO:0016337 | single organismal cell-cell adhesion | 5.43E-04 | 3.69E-03 |
| GO:0001664 | G-protein coupled receptor binding | 6.02E-04 | 3.88E-03 |
| GO:0004499 | N,N-dimethylaniline monooxygenase activity | 6.02E-04 | 3.88E-03 |
| GO:0016820 | hydrolase activity, acting on acid anhydrides, catalyzing transmembrane movement of substances | 6.02E-04 | 3.88E-03 |
| GO:0020037 | heme binding | 6.03E-04 | 3.88E-03 |
| GO:0031110 | regulation of microtubule polymerization or depolymerization | 6.02E-04 | 3.88E-03 |
| GO:0006414 | translational elongation | 7.81E-04 | 4.97E-03 |
| GO:0006457 | protein folding | 8.67E-04 | 5.46E-03 |
| GO:0015934 | large ribosomal subunit | 9.08E-04 | 5.66E-03 |
| GO:0004871 | signal transducer activity | 1.17E-03 | 7.25E-03 |
| GO:0005328 | neurotransmitter:sodium symporter activity | 1.31E-03 | 7.93E-03 |
| GO:0040007 | growth | 1.31E-03 | 7.93E-03 |
| GO:0005200 | structural constituent of cytoskeleton | 1.41E-03 | 8.43E-03 |
| GO:0006955 | immune response | 1.64E-03 | 9.74E-03 |
| GO:0004252 | serine-type endopeptidase activity | 1.80E-03 | 1.06E-02 |
| GO:0004427 | inorganic diphosphatase activity | 2.24E-03 | 1.12E-02 |
| GO:0004618 | phosphoglycerate kinase activity | 2.24E-03 | 1.12E-02 |

| | | | |
|---|---|---|---|
| GO:0004726 | non-membrane-spanning protein tyrosine phosphatase activity | 2.24E-03 | 1.12E-02 |
| GO:0004800 | thyroxine 5'-deiodinase activity | 2.24E-03 | 1.12E-02 |
| GO:0005094 | Rho GDP-dissociation inhibitor activity | 2.24E-03 | 1.12E-02 |
| GO:0005158 | insulin receptor binding | 2.21E-03 | 1.12E-02 |
| GO:0006796 | phosphate-containing compound metabolic process | 2.24E-03 | 1.12E-02 |
| GO:0007269 | neurotransmitter secretion | 2.24E-03 | 1.12E-02 |
| GO:0007420 | brain development | 2.24E-03 | 1.12E-02 |
| GO:0015016 | [heparan sulfate]-glucosamine N-sulfotransferase activity | 2.24E-03 | 1.12E-02 |
| GO:0015074 | DNA integration | 1.96E-03 | 1.12E-02 |
| GO:0016471 | vacuolar proton-transporting V-type ATPase complex | 2.24E-03 | 1.12E-02 |
| GO:0016594 | glycine binding | 2.24E-03 | 1.12E-02 |
| GO:0016934 | extracellular-glycine-gated chloride channel activity | 2.24E-03 | 1.12E-02 |
| GO:0022824 | transmitter-gated ion channel activity | 2.24E-03 | 1.12E-02 |
| GO:0032968 | positive regulation of transcription elongation from RNA polymerase II promoter | 2.24E-03 | 1.12E-02 |
| GO:0035329 | hippo signaling | 2.24E-03 | 1.12E-02 |
| GO:0045263 | proton-transporting ATP synthase complex, coupling factor F(o) | 2.24E-03 | 1.12E-02 |

| | | | |
|---|---|---|---|
| GO:0055085 | transmembrane transport | 2.44E-03 | 1.21E-02 |
| GO:0006950 | response to stress | 3.28E-03 | 1.60E-02 |
| GO:0015992 | proton transport | 3.28E-03 | 1.60E-02 |
| GO:0005216 | ion channel activity | 3.45E-03 | 1.67E-02 |
| GO:0004601 | peroxidase activity | 4.80E-03 | 2.27E-02 |
| GO:0005742 | mitochondrial outer membrane translocase complex | 4.80E-03 | 2.27E-02 |
| GO:0007179 | transforming growth factor beta receptor signaling pathway | 4.80E-03 | 2.27E-02 |
| GO:0006821 | chloride transport | 4.86E-03 | 2.29E-02 |
| GO:0006006 | glucose metabolic process | 6.86E-03 | 3.17E-02 |
| GO:0019773 | proteasome core complex, alpha-subunit complex | 6.86E-03 | 3.17E-02 |
| GO:0005578 | proteinaceous extracellular matrix | 6.94E-03 | 3.19E-02 |
| GO:0006836 | neurotransmitter transport | 7.05E-03 | 3.21E-02 |
| GO:0006413 | translational initiation | 7.62E-03 | 3.42E-02 |
| GO:0030286 | dynein complex | 7.62E-03 | 3.42E-02 |
| GO:0002224 | toll-like receptor signaling pathway | 1.03E-02 | 3.81E-02 |
| GO:0004146 | dihydrofolate reductase activity | 1.03E-02 | 3.81E-02 |
| GO:0004356 | glutamate-ammonia ligase activity | 1.03E-02 | 3.81E-02 |
| GO:0004392 | heme oxygenase (decyclizing) activity | 1.03E-02 | 3.81E-02 |
| GO:0004563 | beta-N-acetylhexosaminidase activity | 1.03E-02 | 3.81E-02 |

| | | | |
|---|---|---|---|
| GO:0004656 | procollagen-proline 4-dioxygenase activity | 1.03E-02 | 3.81E-02 |
| GO:0005030 | neurotrophin receptor activity | 1.03E-02 | 3.81E-02 |
| GO:0005853 | eukaryotic translation elongation factor 1 complex | 1.03E-02 | 3.81E-02 |
| GO:0005960 | glycine cleavage complex | 9.24E-03 | 3.81E-02 |
| GO:0006471 | protein ADP-ribosylation | 9.80E-03 | 3.81E-02 |
| GO:0006542 | glutamine biosynthetic process | 1.03E-02 | 3.81E-02 |
| GO:0006545 | glycine biosynthetic process | 1.03E-02 | 3.81E-02 |
| GO:0006788 | heme oxidation | 1.03E-02 | 3.81E-02 |
| GO:0006885 | regulation of pH | 9.80E-03 | 3.81E-02 |
| GO:0007195 | adenylate cyclase-inhibiting dopamine receptor signaling pathway | 1.03E-02 | 3.81E-02 |
| GO:0009263 | deoxyribonucleotide biosynthetic process | 1.03E-02 | 3.81E-02 |
| GO:0009408 | response to heat | 9.24E-03 | 3.81E-02 |
| GO:0015321 | sodium-dependent phosphate transmembrane transporter activity | 1.03E-02 | 3.81E-02 |
| GO:0015385 | sodium:proton antiporter activity | 9.80E-03 | 3.81E-02 |
| GO:0016494 | C-X-C chemokine receptor activity | 1.03E-02 | 3.81E-02 |
| GO:0016607 | nuclear speck | 1.03E-02 | 3.81E-02 |
| GO:0019464 | glycine decarboxylation via glycine cleavage system | 9.24E-03 | 3.81E-02 |
| GO:0019538 | protein metabolic process | 1.03E-02 | 3.81E-02 |

| | | | |
|---|---|---|---|
| GO:0031032 | actomyosin structure organization | 9.24E-03 | 3.81E-02 |
| GO:0031072 | heat shock protein binding | 9.80E-03 | 3.81E-02 |
| GO:0043560 | insulin receptor substrate binding | 1.03E-02 | 3.81E-02 |
| GO:0044341 | sodium-dependent phosphate transport | 1.03E-02 | 3.81E-02 |
| GO:0046541 | saliva secretion | 1.03E-02 | 3.81E-02 |
| GO:0070461 | SAGA-type complex | 1.03E-02 | 3.81E-02 |
| GO:0051015 | actin filament binding | 1.17E-02 | 4.31E-02 |

**Table S16.** Functionally enriched KEGG pathway categories of sika deer contracted genes.

| ID | Categories | *P*-value | *Padj* |
|---|---|---|---|
| ko04030 | G protein-coupled receptors | 0 | 0 |
| ko04740 | Olfactory transduction | 0 | 0 |
| ko04040 | Ion channels | 3.24E-11 | 3.52E-09 |
| ko00590 | Arachidonic acid metabolism | 5.27E-11 | 4.29E-09 |
| ko05020 | Prion diseases | 3.15E-08 | 2.05E-06 |
| ko02000 | Transporters | 1.29E-07 | 6.99E-06 |
| ko04330 | Notch signaling pathway | 1.59E-07 | 7.43E-06 |
| ko04975 | Fat digestion and absorption | 2.20E-06 | 8.95E-05 |
| ko00591 | Linoleic acid metabolism | 3.46E-06 | 1.13E-04 |
| ko04516 | Cell adhesion molecules and their ligands | 3.16E-06 | 1.13E-04 |
| ko04020 | Calcium signaling pathway | 1.57E-05 | 4.66E-04 |
| ko00199 | Cytochrome P450 | 3.40E-05 | 7.91E-04 |
| ko04320 | Dorso-ventral axis formation | 3.36E-05 | 7.91E-04 |
| ko04960 | Aldosterone-regulated sodium reabsorption | 3.28E-05 | 7.91E-04 |
| ko04668 | TNF signaling pathway | 5.74E-05 | 1.25E-03 |
| ko00592 | alpha-Linolenic acid metabolism | 6.93E-05 | 1.34E-03 |
| ko05146 | Amoebiasis | 7.01E-05 | 1.34E-03 |
| ko03320 | PPAR signaling pathway | 7.50E-05 | 1.36E-03 |

| | | | |
|---|---|---|---|
| ko04014 | Ras signaling pathway | 1.25E-04 | 2.11E-03 |
| ko04621 | NOD-like receptor signaling pathway | 1.29E-04 | 2.11E-03 |
| ko04978 | Mineral absorption | 1.37E-04 | 2.13E-03 |
| ko04973 | Carbohydrate digestion and absorption | 1.57E-04 | 2.32E-03 |
| ko05164 | Influenza A | 1.73E-04 | 2.45E-03 |
| ko04052 | Cytokines | 2.28E-04 | 3.10E-03 |
| ko05222 | Small cell lung cancer | 2.65E-04 | 3.45E-03 |
| ko04972 | Pancreatic secretion | 2.92E-04 | 3.66E-03 |
| ko04380 | Osteoclast differentiation | 3.53E-04 | 4.26E-03 |
| ko04360 | Axon guidance | 3.78E-04 | 4.41E-03 |
| ko05134 | Legionellosis | 3.96E-04 | 4.45E-03 |
| ko04010 | MAPK signaling pathway | 4.55E-04 | 4.95E-03 |
| ko04622 | RIG-I-like receptor signaling pathway | 5.43E-04 | 5.53E-03 |
| ko04658 | Th1 and Th2 cell differentiation | 5.28E-04 | 5.53E-03 |
| ko00062 | Fatty acid elongation | 6.07E-04 | 5.82E-03 |
| ko04657 | IL-17 signaling pathway | 5.91E-04 | 5.82E-03 |
| ko00565 | Ether lipid metabolism | 6.94E-04 | 6.46E-03 |
| ko04660 | T cell receptor signaling pathway | 7.36E-04 | 6.66E-03 |
| ko04742 | Taste transduction | 8.57E-04 | 7.55E-03 |
| ko04210 | Apoptosis | 1.19E-03 | 1.02E-02 |

| | | | |
|---|---|---|---|
| ko04726 | Serotonergic synapse | 1.29E-03 | 1.08E-02 |
| ko04066 | HIF-1 signaling pathway | 1.68E-03 | 1.34E-02 |
| ko05152 | Tuberculosis | 1.68E-03 | 1.34E-02 |
| ko05167 | Kaposi's sarcoma-associated herpesvirus infection | 2.24E-03 | 1.74E-02 |
| ko04662 | B cell receptor signaling pathway | 2.69E-03 | 2.03E-02 |
| ko05165 | Human papillomavirus infection | 2.74E-03 | 2.03E-02 |
| ko04750 | Inflammatory mediator regulation of TRP channels | 3.11E-03 | 2.26E-02 |
| ko04121 | Ubiquitin system | 3.49E-03 | 2.46E-02 |
| ko04217 | Necroptosis | 3.55E-03 | 2.46E-02 |
| ko04919 | Thyroid hormone signaling pathway | 3.72E-03 | 2.50E-02 |
| ko04930 | Type II diabetes mellitus | 3.75E-03 | 2.50E-02 |
| ko01522 | Endocrine resistance | 4.86E-03 | 3.11E-02 |
| ko04620 | Toll-like receptor signaling pathway | 4.86E-03 | 3.11E-02 |
| ko05160 | Hepatitis C | 5.48E-03 | 3.37E-02 |
| ko99992 | Membrane and intracellular structural | 5.41E-03 | 3.37E-02 |
| ko04971 | Gastric acid secretion | 5.69E-03 | 3.44E-02 |
| ko05211 | Renal cell carcinoma | 7.40E-03 | 4.39E-02 |
| ko04911 | Insulin secretion | 8.33E-03 | 4.76E-02 |
| ko04925 | Aldosterone synthesis and secretion | 8.33E-03 | 4.76E-02 |

**Table S17.** Functionally enriched GO categories of sika deer contracted genes.

| ID | Categories | *P*-value | *Padj* |
|---|---|---|---|
| GO:0004930 | G-protein coupled receptor activity | 0 | 0 |
| GO:0004984 | olfactory receptor activity | 0 | 0 |
| GO:0005509 | calcium ion binding | 0 | 0 |
| GO:0005515 | protein binding | 0 | 0 |
| GO:0007186 | G-protein coupled receptor signaling pathway | 0 | 0 |
| GO:0006820 | anion transport | 2.21E-12 | 1.45E-10 |
| GO:0016021 | integral component of membrane | 3.23E-12 | 1.82E-10 |
| GO:0005506 | iron ion binding | 4.97E-12 | 2.45E-10 |
| GO:0005615 | extracellular space | 3.30E-10 | 1.44E-08 |
| GO:0008272 | sulfate transport | 4.48E-10 | 1.60E-08 |
| GO:0015116 | sulfate transmembrane transporter activity | 4.48E-10 | 1.60E-08 |
| GO:0005452 | inorganic anion exchanger activity | 6.21E-09 | 2.04E-07 |
| GO:0022857 | transmembrane transporter activity | 7.14E-09 | 2.16E-07 |
| GO:0008271 | secondary active sulfate transmembrane transporter activity | 1.97E-08 | 5.54E-07 |
| GO:0005215 | transporter activity | 2.88E-08 | 7.56E-07 |
| GO:0016705 | oxidoreductase activity, acting on paired donors, with incorporation or reduction of molecular oxygen | 3.35E-08 | 8.25E-07 |

| | | | |
|---|---|---|---|
| GO:0005149 | interleukin-1 receptor binding | 4.38E-08 | 9.62E-07 |
| GO:0055085 | transmembrane transport | 4.39E-08 | 9.62E-07 |
| GO:0008509 | anion transmembrane transporter activity | 8.40E-08 | 1.74E-06 |
| GO:0005245 | voltage-gated calcium channel activity | 1.43E-07 | 2.61E-06 |
| GO:0005891 | voltage-gated calcium channel complex | 1.43E-07 | 2.61E-06 |
| GO:0016020 | membrane | 1.46E-07 | 2.61E-06 |
| GO:0070588 | calcium ion transmembrane transport | 3.01E-07 | 5.15E-06 |
| GO:0005391 | sodium:potassium-exchanging ATPase activity | 7.40E-07 | 1.17E-05 |
| GO:0030212 | hyaluronan metabolic process | 7.40E-07 | 1.17E-05 |
| GO:0006810 | transport | 2.57E-06 | 3.89E-05 |
| GO:0004623 | phospholipase A2 activity | 3.24E-06 | 4.40E-05 |
| GO:0007219 | Notch signaling pathway | 3.24E-06 | 4.40E-05 |
| GO:0050482 | arachidonic acid secretion | 3.24E-06 | 4.40E-05 |
| GO:0006814 | sodium ion transport | 6.64E-06 | 8.73E-05 |
| GO:0005576 | extracellular region | 7.95E-06 | 1.01E-04 |
| GO:0005216 | ion channel activity | 1.08E-05 | 1.27E-04 |
| GO:0006644 | phospholipid metabolic process | 1.09E-05 | 1.27E-04 |
| GO:0016702 | oxidoreductase activity, acting on single donors with incorporation of molecular oxygen, incorporation of two atoms of oxygen | 1.09E-05 | 1.27E-04 |

| | | | |
|---|---|---|---|
| GO:0006596 | polyamine biosynthetic process | 1.25E-05 | 1.40E-04 |
| GO:0016042 | lipid catabolic process | 1.84E-05 | 2.02E-04 |
| GO:0006952 | defense response | 2.23E-05 | 2.38E-04 |
| GO:0006811 | ion transport | 3.15E-05 | 3.26E-04 |
| GO:0020037 | heme binding | 3.64E-05 | 3.67E-04 |
| GO:0005579 | membrane attack complex | 5.94E-05 | 5.20E-04 |
| GO:0006406 | mRNA export from nucleus | 5.94E-05 | 5.20E-04 |
| GO:0006839 | mitochondrial transport | 5.94E-05 | 5.20E-04 |
| GO:0009922 | fatty acid elongase activity | 5.94E-05 | 5.20E-04 |
| GO:0019367 | fatty acid elongation, saturated fatty acid | 5.94E-05 | 5.20E-04 |
| GO:0042761 | very long-chain fatty acid biosynthetic process | 5.94E-05 | 5.20E-04 |
| GO:0004190 | aspartic-type endopeptidase activity | 9.14E-05 | 7.66E-04 |
| GO:0007156 | homophilic cell adhesion via plasma membrane adhesion molecules | 9.07E-05 | 7.66E-04 |
| GO:0004197 | cysteine-type endopeptidase activity | 1.38E-04 | 1.13E-03 |
| GO:0008289 | lipid binding | 1.54E-04 | 1.24E-03 |
| GO:0004176 | ATP-dependent peptidase activity | 1.70E-04 | 1.31E-03 |
| GO:0030154 | cell differentiation | 1.70E-04 | 1.31E-03 |
| GO:0001733 | galactosylceramide sulfotransferase activity | 2.10E-04 | 1.40E-03 |
| GO:0005219 | ryanodine-sensitive calcium-release channel activity | 2.10E-04 | 1.40E-03 |

| | | | |
|---|---|---|---|
| GO:0006874 | cellular calcium ion homeostasis | 2.10E-04 | 1.40E-03 |
| GO:0008191 | metalloendopeptidase inhibitor activity | 2.10E-04 | 1.40E-03 |
| GO:0008308 | voltage-gated anion channel activity | 2.10E-04 | 1.40E-03 |
| GO:0009247 | glycolipid biosynthetic process | 2.10E-04 | 1.40E-03 |
| GO:0030866 | cortical actin cytoskeleton organization | 2.10E-04 | 1.40E-03 |
| GO:0044070 | regulation of anion transport | 2.10E-04 | 1.40E-03 |
| GO:0016817 | hydrolase activity, acting on acid anhydrides | 2.52E-04 | 1.66E-03 |
| GO:0005886 | plasma membrane | 2.64E-04 | 1.71E-03 |
| GO:0007166 | cell surface receptor signaling pathway | 2.95E-04 | 1.87E-03 |
| GO:0005337 | nucleoside transmembrane transporter activity | 3.77E-04 | 2.25E-03 |
| GO:0015280 | ligand-gated sodium channel activity | 3.77E-04 | 2.25E-03 |
| GO:0031966 | mitochondrial membrane | 3.77E-04 | 2.25E-03 |
| GO:0051262 | protein tetramerization | 3.77E-04 | 2.25E-03 |
| GO:0071805 | potassium ion transmembrane transport | 4.37E-04 | 2.57E-03 |
| GO:0007155 | cell adhesion | 4.53E-04 | 2.62E-03 |
| GO:0004888 | transmembrane signaling receptor activity | 6.12E-04 | 3.49E-03 |
| GO:0004866 | endopeptidase inhibitor activity | 6.36E-04 | 3.58E-03 |
| GO:0004842 | ubiquitin-protein transferase activity | 7.32E-04 | 4.06E-03 |
| GO:0004683 | calmodulin-dependent protein kinase activity | 8.03E-04 | 4.39E-03 |
| GO:0005267 | potassium channel activity | 1.10E-03 | 5.91E-03 |

| | | | |
|---|---|---|---|
| GO:0005272 | sodium channel activity | 1.23E-03 | 6.48E-03 |
| GO:0006887 | exocytosis | 1.23E-03 | 6.48E-03 |
| GO:0008234 | cysteine-type peptidase activity | 1.57E-03 | 8.17E-03 |
| GO:0016810 | hydrolase activity, acting on carbon-nitrogen (but not peptide) bonds | 1.86E-03 | 9.39E-03 |
| GO:0031418 | L-ascorbic acid binding | 1.86E-03 | 9.39E-03 |
| GO:0008023 | transcription elongation factor complex | 1.92E-03 | 9.57E-03 |
| GO:0000145 | exocyst | 1.96E-03 | 9.65E-03 |
| GO:0030414 | peptidase inhibitor activity | 2.93E-03 | 1.43E-02 |
| GO:0004867 | serine-type endopeptidase inhibitor activity | 3.38E-03 | 1.53E-02 |
| GO:0006935 | chemotaxis | 3.38E-03 | 1.53E-02 |
| GO:0008009 | chemokine activity | 3.38E-03 | 1.53E-02 |
| GO:0008158 | hedgehog receptor activity | 3.54E-03 | 1.53E-02 |
| GO:0015276 | ligand-gated ion channel activity | 3.54E-03 | 1.53E-02 |
| GO:0031047 | gene silencing by RNA | 3.54E-03 | 1.53E-02 |
| GO:0046373 | L-arabinose metabolic process | 3.54E-03 | 1.53E-02 |
| GO:0046556 | alpha-L-arabinofuranosidase activity | 3.54E-03 | 1.53E-02 |
| GO:0050793 | regulation of developmental process | 3.54E-03 | 1.53E-02 |
| GO:0051028 | mRNA transport | 3.54E-03 | 1.53E-02 |
| GO:0006368 | transcription elongation from RNA polymerase II | 3.67E-03 | 1.55E-02 |

| | | | |
|---|---|---|---|
| | promoter | | |
| GO:0015631 | tubulin binding | 3.67E-03 | 1.55E-02 |
| GO:0005741 | mitochondrial outer membrane | 5.78E-03 | 2.42E-02 |
| GO:0042981 | regulation of apoptotic process | 6.06E-03 | 2.51E-02 |
| GO:0030695 | GTPase regulator activity | 6.13E-03 | 2.52E-02 |
| GO:0006508 | proteolysis | 6.31E-03 | 2.56E-02 |
| GO:0003779 | actin binding | 7.58E-03 | 3.05E-02 |
| GO:0004872 | receptor activity | 8.60E-03 | 3.41E-02 |
| GO:0007275 | multicellular organismal development | 8.65E-03 | 3.41E-02 |
| GO:0004013 | adenosylhomocysteinase activity | 1.02E-02 | 3.72E-02 |
| GO:0004332 | fructose-bisphosphate aldolase activity | 1.02E-02 | 3.72E-02 |
| GO:0004645 | phosphorylase activity | 1.02E-02 | 3.72E-02 |
| GO:0004887 | thyroid hormone receptor activity | 1.02E-02 | 3.72E-02 |
| GO:0008140 | cAMP response element binding protein binding | 1.02E-02 | 3.72E-02 |
| GO:0008184 | glycogen phosphorylase activity | 1.02E-02 | 3.72E-02 |
| GO:0019510 | S-adenosylhomocysteine catabolic process | 1.02E-02 | 3.72E-02 |
| GO:0032947 | protein complex scaffold | 1.02E-02 | 3.72E-02 |
| GO:0005634 | nucleus | 1.06E-02 | 3.78E-02 |
| GO:0008270 | zinc ion binding | 1.05E-02 | 3.78E-02 |
| GO:0004725 | protein tyrosine phosphatase activity | 1.09E-02 | 3.86E-02 |

| | | | |
|---|---|---|---|
| GO:0007165 | signal transduction | 1.17E-02 | 4.13E-02 |
| GO:0006816 | calcium ion transport | 1.22E-02 | 4.21E-02 |
| GO:0016567 | protein ubiquitination | 1.22E-02 | 4.21E-02 |
| GO:0006955 | immune response | 1.28E-02 | 4.37E-02 |
| GO:0015671 | oxygen transport | 1.35E-02 | 4.57E-02 |
| GO:0006334 | nucleosome assembly | 1.40E-02 | 4.69E-02 |
| GO:0016311 | dephosphorylation | 1.40E-02 | 4.69E-02 |

**Table S18.** Numbers of annotated *UGT* genes in 19 species.

| | ***UGT 2B*** | ***UGT 2C*** | ***UGT 2A3*** | ***UGT 2A1*** | ***UGT 1A*** | ***UGT 8*** | ***UGT 3A*** | ***Total*** |
|---|---|---|---|---|---|---|---|---|
| *Human* | 7 | 0 | 1 | 1 | 1 | 1 | 2 | 13 |
| *Mouse* | 7 | 0 | 1 | 1 | 1 | 1 | 2 | 13 |
| *Single-humped camel* | 3 | 2 | 0 | 1 | 1 | 1 | 1 | 9 |
| *Double-humped camel* | 1 | 1 | 0 | 1 | 1 | 1 | 1 | 6 |
| *Pig* | 4 | 1 | 1 | 1 | 1 | 0 | 1 | 9 |
| *Minke whale* | 1 | 1 | 1 | 0 | 1 | 1 | 1 | 6 |
| *Okapi* | 4 | 5 | 1 | 0 | 1 | 1 | 1 | 13 |
| *Giraffe* | 6 | 2 | 0 | 0 | 1 | 1 | 1 | 11 |
| *Musk deer* | 3 | 2 | 0 | 0 | 3 | 1 | 1 | 10 |
| *Cattle* | 4 | 3 | 0 | 1 | 1 | 1 | 2 | 12 |
| *Yak* | 6 | 5 | 1 | 0 | 1 | 1 | 1 | 15 |
| *Goat* | 7 | 4 | 0 | 1 | 1 | 1 | 2 | 16 |
| *Sheep* | 5 | 3 | 1 | 0 | 1 | 1 | 2 | 13 |
| *Roe deer* | 4 | 3 | 0 | 0 | 1 | 1 | 1 | 10 |
| *Reindeer* | 8 | 3 | 1 | 0 | 1 | 0 | 1 | 14 |
| *White-tailed deer* | 6 | 4 | 0 | 1 | 5 | 1 | 2 | 19 |
| *Milu* | 6 | 4 | 1 | 0 | 2 | 1 | 2 | 16 |

| | | | | | | | | |
|---|---|---|---|---|---|---|---|---|
| *Red deer* | 13 | 6 | 3 | 0 | 1 | 0 | 2 | 25 |
| *Sika deer* | 15 | 5 | 1 | 2 | 2 | 0 | 2 | 27 |

**Table S19.** Design of the feeding experiment.

| | 0 MOL<br><br>(y0 group) | 50% MOL<br><br>(y50 group) | 100% MOL<br><br>(y100 group) | 100% MOL<br><br>(m100 group) |
|---|---|---|---|---|
| Sika deer | 15 tissues*3 | 15 tissues*3 | 15 tissues*3 | 15 tissues*3 |
| | 0 GA<br><br>(n group) | | 10% GA<br><br>(y group) | |
| Cattle | 8 tissues*3[8] | | 8 tissues*3 | |

Twelve sika deer were divided into four groups. Six cattle were divided into two groups.

[8]Two samples were not sequenced due to the pool quality.
